# A roadmap for fair reuse of public microbiome data

**DOI:** 10.1101/2024.06.21.599698

**Authors:** Laura A. Hug, Roland Hatzenpichler, Cristina Moraru, André Soares, Folker Meyer, Anke Heyder, The Data Reuse Consortium, Alexander J. Probst

**Author notes:** For a full list of contributors and affiliations of members of The Data Reuse Consortium please see Supplementary Material. The Data Reuse Core Team; these authors contributed equally to the manuscript.

## Abstract

Science benefits from rapid, open data sharing but samples for sequencing data are expensive for data creators to acquire and process. Current guidelines for data reuse were established two decades ago, when databases were several million times smaller, necessitating an update. This article presents a roadmap to establish best practices for sequence data reuse, developed in consultation with a data consortium of 167 microbiome scientists. It introduces a Data Reuse Information tag (DRI) for public sequencing data, which will be associated with at least one Open Researcher and Contributor ID (ORCID) account. The machine-readable DRI tag indicates that the data creators prefer to be contacted prior to data reuse, and simultaneously provides data consumers with a mechanism to get in touch with the data creators. Ideally, the DRI will facilitate and foster collaborations, and serve as a guideline that can be expanded to other data types.

## Main

Sequencing data reuse has been an evolving topic over the past two decades. The Fort Lauderdale Agreement (FLA) was coined in 2003, prior to the advent of metagenomics and during a time when sequencing was still too costly to be performed by individual laboratories^1^. The FLA concluded that large genome projects should be released before publication in order to accelerate the advancement of science. The FLA strengthened the Bermuda Principles defined in 1996^1^, which advocated the release of sequence data 24 hours after generation and prior to publication of research papers. In 2009, after the Human Genome Project highlighted the advantages of sharing data early and widely, the Toronto Statement (TOR)^2^ brought further attention to pre-publication data sharing. Finally, in 2014, 141 UN member states and the European Union entered into the Nagoya Protocol (https://www.cbd.int/abs/nagoya-protocol/signatories/; Regulation (EU) No 511/2014), which calls on data creators and data users to develop, update and use voluntary codes of conduct, guidelines and best practices in relation to access and benefit-sharing of genetic data (see Box 1 for descriptions of the different roles of researchers working with sequence datasets). Large-scale sequence data analysis has become mainstream with a wide array of tools available, making data mining accessible to many labs. Now, ∼20 years after the FLA, GenBank holds an estimated ∼2.57 Tbp (02/2024) of biological sequencing data, and the Sequence Read Archive (SRA) holds 90.89 Pbp as of February 2024 (***Figure S1***, for details on repositories and abbreviations, please see ***Box 2***). These databases are several million times larger than the available sequence data at the time that the FLA or TOR were formulated. With the rapid, continuous increase in public sequencing data (projected to reach ∼500 Pbp in 2030, ***Figure S1***), data mining projects (or those requiring large public datasets for AI training) have increased in both frequency and scope, necessitating a revisit and potential overhaul of the 20-year old guidelines depicted in the FLA and TOR^1,2^.

Biological sciences, and particularly its subdisciplines associated with generating sequencing data, have been at the forefront of data availability compared to the fields of earth sciences, mathematics, physics, and chemistry^3^. For instance, while open data and data sharing have been practiced in the field of physics education for some time, a formal prospective system to facilitate open data and data mining was only introduced in 2021^4^. In physics, multi-user facilities advocate for data sharing rules to facilitate data mining by scientists who were not part of the original experiments^5^. By contrast, astrophysics is a subdiscipline that traditionally relies on data sharing and reuse due to the exorbitant costs of research data. A study investigating the motivational factors behind data sharing and reuse within astrophysics identified several demotivating factors^5^. Among them were the lack of data standards, the lack of facilitating platforms, inconsistency between datasets, limited documentation, difficulties finding data, and reuse data, and last but not least, competition and fear of being “scooped” (accidentally or purposefully)^5^. The latter point is also considered in the FLA. While the FLA recommends swift pre-publication of data generated by large sequencing consortia, it also states that “[…] *the contribution and interests of the large-scale* data creators [*data producers*] *should be recognized and respected by the users of the data, and the ability of the production centers to analyze and publish their own data should be supported by their funding agencies* […]”. This highlights one of the significant and enduring tensions between data creators and data consumers. Both data creators and data consumers are indispensable to advancing biological sciences, particularly in the realm of sequence data analysis, where many data creators also act as data consumers. Unrestricted public use of microbiome data, on which data creators have not yet published, does not always align with the interests of data creators.

How to achieve unrestricted data usage and, at the same time, give due credit to data creators has been much discussed by the scientific community^6^. In this spirit, a recent study, which thoroughly analyzed the pros and cons of early data release and considered both the needs of data creators and consumers, proposed immediate, unrestricted release of (unpublished upon) sequencing data, in parallel with the adoption of a reward system (*e.g.*, separate promotion and tenure tracks) that acknowledges data creators by universities and research institutions^7^. In addition, the authors proposed making the datasets and the protocols used for their generation citable through Digital Object Identifiers (DOIs). If implemented, these measures would create a safer environment for data sharing, benefiting all parties involved and most importantly, supporting the advancement of science.

Mechanisms for crediting data creators beyond citing associated publications have not yet been implemented in the scientific community. Creating separate tenure tracks or other incentives for data creators and data consumers requires sizable changes in evaluation criteria and would require substantial time to propagate through institutions. DOIs for datasets, on the other hand, seem relatively easy to implement and would provide data creators with a reportable impact metric. However, their use has not yet been widely adopted, possibly due to associated costs with purchasing and maintaining DOIs, which can be prohibitive for many publicly funded research institutions. Potential measures to lower DOI costs would include large-scale agreements between research institutions and DOI providers. Other mechanisms of data citation have been discussed in the community but have not been implemented^8,9^. Currently, data creators do not have any incentive nor reward for releasing sequencing data prior to an associated publication.

It is crucial to implement methodological and ethical guidelines that are based on the principles of good scientific practice, and which are driven by the scientific community to facilitate appropriate use of public data. This need has been highlighted by recent conflicts between data creators and data consumers that played out over social media. Implementing and following guidelines for unpublished data usage by all scientists would create “safe spaces” for data creators to publish their first analyses of data - particularly if they are delayed by resource, time, or personnel constraints. The research topic also affects the expectations for open data. Research related to public health necessitates swift data release to counteract pandemics or identify zoonotic diseases. For example, in the event of a pandemic, there should be no data restriction on research related to the pandemic^10^. The general goal should be to promote open sharing of complete datasets as early and as widely as possible, across all institutions and individuals, which necessitates a technical framework that enhances the communication between data creators and data consumers regarding data reuse.

Here, we propose a roadmap to enable fair use of public microbiome data (***Figure 1***). This roadmap (1) addresses the lack of consensus in the field of microbiome research regarding public microbiome data use and reuse, (2) promotes communication between data consumers and data creators, and (3) facilitates the rapid advancement of the microbiome field, including supporting continued increases in data mining. This roadmap and its adoption by the scientific community (155 scientists as part of the Data Reuse Consortium - see ***Table S1*** - totaling 162 supporters, including the co-authors of this paper) will provide a citable resource regarding “best practices for public data reuse”, will enable appropriate data reuse by data consumers, and will reduce tension for data creators when submitting data.

**Figure 1:**
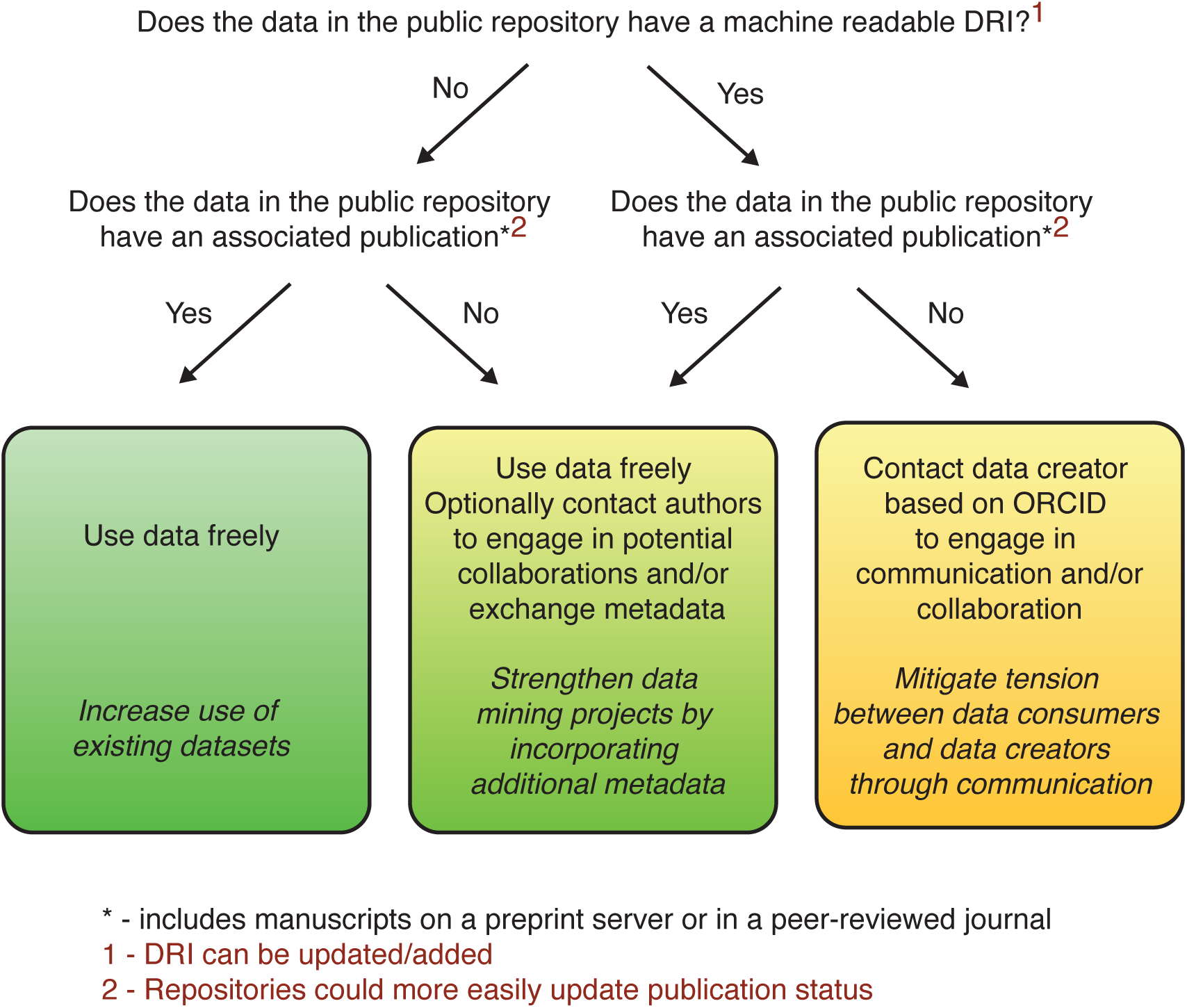
Recommendations for best practice for fair reuse of public microbiome data applicable unless there is a differing policy in place by the institution employing the data consumer or there is a restricting license associated with the dataset itself.

We assessed community interest in and likelihood of adopting the proposed roadmap through an anonymous survey that gathered responses from 306 scientists representing all continents (except Antarctica; ***Figure S2*, *Table S1***). The roadmap was refined through conversations with Data Reuse Consortium members, the Genome Reference Consortium, and other entities involved in sequence data management (*e.g.*, the JGI).

To achieve the goal of this roadmap, we propose the introduction of a new machine-readable metadata tag, named Data Reuse Information (DRI), containing ORCIDs of the data creators associated with data in public databases. The DRI will clearly indicate the point of contact for communication and that communication is desired by data creators. Ideally, the ability to provide a point of contact for data reuse will lead to more rapid and complete data deposition. Following adoption by databases, authors and scientific journals would ideally integrate statements confirming that the best practices governed by DRI use were used in manuscripts and submission processes. Ultimately, this roadmap outlines the expected practices for open data use for sequencing data and represents a model for other biological data such as metabolomics or proteomics data.

### Box 1

#### Definitions of roles of scientists/legal entities related to microbiome and sequence data

**Data consumer**: Any legal entity interested in using public data.

**Data creator**: Entity(/ies), *i.e.*, individual or multiple researchers, who designed the study, obtained the samples, and intended to publish an analysis; typically assumed to have priority on analysis.

**Data distributor**: Public databases that provide access to the digital data (*e.g.*, GenBank, ENA, DDBJ, please see ***Box 2***).

**Data generator**: Entity that renders the sample into a digital object (*e.g.*, a sequencing facility that processes biological material and produces FASTQ and other sequence-related files to transmit to another entity).

**Data owner**: Legal entity who owns the data by rights. This can be different from “data creator” (*e.g.*, an institute at which the data creator is employed or a nation/state.)

### Box 2

#### Abbreviations extensively used in microbiome data research

**COGs - Clusters of Orthologous Groups.** Represents a collection of proteins from complete bacterial and archaeal genomes, grouped into clusters of orthologs, and associated with functional annotations.

**DDBJ - DNA Data Bank of Japan**. A database of nucleotide sequence data maintained by the National Institute of Genetics (NIG) in Japan.

**EMBL - European Molecular Biology Laboratory.** A research organization that conducts basic research in molecular biology and offers a range of scientific resources..

**EMBL-EBI - European Bioinformatics Institute.** A bioinformatics research center belonging to EMBL, which maintains and provides access to several sequence-related databases (e.g., ENA, Interpro, PDBe, UniProt).

**ENA - European Nucleotide Archive.** A database of raw sequencing data and annotated sequence data, from a wide range of organisms, maintained by EMBL-EBI.

**GenBank** - **Genetic Sequence Database**. A database of DNA and RNA sequences from a wide range of organisms, along with associated annotations and metadata.

**GSC - Genomic Standards Consortium.** An international organization dedicated to the development and implementation of standards and best practices in genomics and related fields.

**IMG/M - Integrated Microbial Genomes and Metagenomes.** A data management and analysis system for microbial genomes and metagenomes maintained by the Department of Energy’s Joint Genome Institute (JGI) in the USA.

**IMG/VR - Integrated Microbial Genomes with Virus-Related datasets.** A specialized database storing virus genomic and metagenomic sequences, annotations and metadata.

**InterPro - Integrated resource of protein domains and functional sites.** A database that integrates information on protein domains, motifs, and functional sites from a variety of sources.

**INSDC - International Nucleotide Sequence Database Collaboration.** A data sharing initiative between DDBJ, EMBL-EBI, and NCBI.

**KEGG - Kyoto Encyclopedia of Genes and Genomes.** A comprehensive database and knowledge base of biological systems, including genes, proteins, and biochemical pathways. It is maintained by Kanehisa Laboratories in Japan.

**KOG - euKaryotic Orthologous Groups.** A collection of proteins from eukaryotic genomes, grouped into clusters of orthologs, and associated with functional annotations.

**NCBI - National Center for Biotechnology Information.** Part of NIH; a central repository for molecular sequence data, including several databases (e.g., GenBank, RefSeq, SRA, COG, KOG, etc.)

**NIH - National Institutes of Health.** A biomedical research agency of the federal government of the USA.

**PDBe - Protein Data Bank in Europe.** A comprehensive collection of 3D structures of proteins and other macromolecules.

**PFAM - Protein family database.** A database of protein families, domains, and functional sites.

**RefSeq - Reference Sequence Database.** A comprehensive, non-redundant database of reference genomic, transcriptomic and proteomic sequences, for a wide range of organisms.

**SRA - Sequence Read Archive.** A public repository for raw sequencing data generated by platforms such as Sanger, Illumina, Ion Torrent, and Pacific Biosciences.

**UniProt - Universal Protein Resource.** A comprehensive protein sequence database, including additional field-specific contextual information (as for example protein domain structure, and known interactions).

### Current practices and data policies

The microbiome data types most frequently reused include amplicon datasets of single genes, as well as reads and assemblies of genomes, and metagenomic and metatranscriptomic reads and assemblies. The sequence data, while typically structured as machine-readable files, come in various formats depending on the hosting repository. Organizations hosting the data typically have their own legal framework governing data download. In the case of, *e.g.*, EMBL-EBI’s ENA database, additional license information can be used to add further restrictions on top of ENA’s base policies. EMBL-EBI’s policy is to use the Creative Commons Zero (CC0) license that, in essence, places the data in the public domain. However, EMBL-EBI clearly states they cannot guarantee that all data in a respective resource are indeed under a CC0 license (https://www.ebi.ac.uk/licencing). This puts the responsibility on the data consumer to ensure their action is indeed legal and covered by the copyrights that may be associated with the data they are accessing, even if the licensing information is not machine-readable. The current state of data policies across online repositories of biological data, including EMBL-EBI, is largely reflective of a now outdated statement in the TOR wherein data users are advised to contact data creators to “discuss publication plans”, a demanding and often, unfeasible approach given today’s rate of data production and availability of new data analysis pipelines. The TOR, from 2009, and the FLA, from 2003, are the agreements guiding the field to date, though their contents are often not well known by microbiome scientists across all career stages (***Figures S4-6***)^1,2^. The FLA specifies, among other things, that “sequence assemblies of 2kb or greater by large-scale sequencing efforts” must be rapidly released. The language around the scale of data shows this comes from a bygone sequencing era, and the emphasis on large-scale data creators is no longer in line with the prevalence of independent labs as data creators today. Interestingly, the conflict between data sharing and publishing a first analysis was acknowledged in the FLA, without providing guidelines for navigating this concern. The TOR elaborated further on the conflict between “data [creators] and data users”, stating that, in the author’s experience, conflicts have rarely arisen. The TOR lists a set of conditions (scale, utility, reference data, community acceptance) to consider for pre-publication data sharing that are of limited relevance for today’s data landscape^2^. Currently, individual research groups can contribute substantial data sets, both in terms of size and scientific value, and the idea of individualized, private agreements on data sharing, as suggested by the TOR, is no longer viable. In response to this need, repositories have developed their own policies (*e.g.*, EMBL-EBI’s license information discussed above). Data distributors recognize that some public data sets, although hosted by their respective services, carry additional restrictions that are currently neither easily visible nor machine readable.

### Identifying conflicts of interest between data consumers and data creators

The current scale of open sequencing data, including massive open datasets such as the *Tara* Oceans and Integrative Human Microbiome projects, has made meta-analyses drawing on public data a powerful avenue to explore microbial systems^11,12^. Use of public data is now routine; close to 80% of respondents to our poll on data usage identified themselves as both data creators and data consumers (***Box 3***, ***Figure 2***). Access to public data has unequivocally improved the depth of the science conducted. However, it is often difficult to assess whether data reuse follows the expectations of communication and collaboration outlined in the TOR. There are many widely used software tools that include use of secondarily accessed data for which no primary publication exists (*e.g.*, the Genome Taxonomy Database, GToTree)^13,14^. Identification of the publication status of data in many repositories is not straightforward. Clarifying and facilitating data reuse is in the best interest of the community. As more governments begin to require data deposition on short or immediate timelines, there is a growing tension between data creators and data consumers around public data use.

The roots of the potential conflict of interest between data creators and data consumers are: (1) a disconnect between the efforts expended by data creators in generating sequence data and associated metadata and the ease of reuse and general lack of acknowledgment of data origins by consumers; (2) shared aims of open science and timely deposition in public databases that contrasts with lengthy timelines for analyses on multi-omic datasets; and (3) the perceived threat that unauthorized data reuse will negatively impact planned research directions, funded research goals, acquisition of new funds, or the career perspectives of early career researchers for both creators and consumers. These are discussed in detail below:

1. Creators sometimes feel that their monetary, time, and intellectual investments to design and conduct sample collection and experiments; obtain permission for, plan, fund, and carry out research expeditions; process samples; and deposit data and metadata are not adequately acknowledged or are potentially ignored by data consumers. Data creators must obtain legal documents (*e.g.*, sampling permits, visas) and follow international agreements (*e.g.*, the Convention on Biological Diversity, Nagoya protocol), and manage the risks that come with certain fieldwork (*e.g.*, treacherous terrain, wilderness areas, areas with high criminal activity). In addition, they must secure funding for custom-design vehicles and instrumentation needed for sample collection (*e.g.*, drill ships, research vessels, submarines, buoys, remote samplers) and maintain research sites in hard-to-access areas (*e.g.*, polar regions or the Amazon). Unbeknownst to data consumers, the original data creators may be bound by restrictive agreements on appropriate or ethical data use if research was conducted in a National Park, on private land, or land owned by Indigenous nations, and/or for samples obtained from human specimens or biobanks^15^. There are currently limited rewards for data creators when their data is reused and data creators have little incentive to make detailed metadata available. Systems for reporting and incentivizing data deposition have been proposed, but are not yet the norm^7,16^.
2. Both data creators and consumers generally share ideals of open science and rapid advancement of science. However, conflicts arise from disagreements in the timing of sharing data and prioritization of access. Data creators must balance long trainee timelines with publication of datasets intended for multiple research questions. Publishing a first paper on a large dataset and depositing the full data may make additional research projects associated with that dataset vulnerable to scooping. Results from our poll suggest that 53% of researchers are concerned about negative impacts on their research program and/or mentees from unauthorized data reuse (***Box 3, Figure 2, Figures S7-S9***). As a result, partial or raw datasets or datasets lacking key metadata are deposited in place of more polished, complete datasets with full metadata to guide interpretation of genome data. In the absence of open data, data consumers are frequently unable to access contextual information (*e.g.*, physicochemical parameters, geolocations) of the field site that are essential to accurately interpret the data, to the detriment of downstream analyses. These issues are exacerbated by public sequence databases lacking (links to) the associated metadata and a general lack of familiarity of many data consumers with the literature on, or the environmental context of, a specific system.
3. Duplication of effort and the potential for lowered impact or difficulties publishing replicated results are a loss for both creators and consumers. For data creators, raw data underlying published scientific results must be made public to meet expectations for reproducibility. However, unrestricted access to public data can compromise permits, site access agreements, and research ethics board approvals, all of which can negatively impact the data creators’ and their mentees’ ongoing research. For data consumers, even unintentional reuse of restricted data can slow research progress while appropriate permissions are sought, delay publications while data is removed, and, in extreme cases, lead to paper retractions. A lack of formal structure for data reuse causes tension for data creators and consumers alike.

### A roadmap to reduce tension between data creators and data consumers

There are multiple potential avenues for mitigating the three conflicts of interest discussed above, yet not every approach is suitable or can be realized. For instance, funding agencies have the power to set rules for data release and data reuse in principle. However, besides the differentiation between private and taxpayer-funded agencies, funders usually have diverging agendas that not only differ across political borders but are also heterogeneous within a single country. To address the current tension(s) between data creators and data consumers and to update the existing agreements from more than 15 years ago, we propose a comprehensive roadmap for data reuse (***Figure 1***). We recommend following this roadmap except in cases in which institutions or funding agencies have a different policy for data reuse in place or there is a restricting license associated with the dataset itself. This roadmap was developed with the aim of minimizing friction between data creators and data consumers and involves the introduction of a new machine-readable DRI metadata tag for facilitating communication between data creators, generators, and consumers.

Transparent, fair and ethical use of public data necessitates clear labeling of its usability by data distributors. Although we initially considered a system that places limitations on free data use, following consultations with the Genomic Standards Consortium and polling 306 microbiome scientists, we have converged on an approach that focuses on both simplicity and openness while achieving nearly all of the desired effects. The DRI tag would attach ORCIDs of the data creators to deposited data, signaling that data creators wish to be contacted (*e.g.*, via email) prior to data use^17^. This way, the DRI tag also provides stable contact information to allow data consumers to easily reach data creators. ORCIDs are both free and ubiquitous and, most importantly, are already used internally by the INSDC community.

The DRI will consist of a tag (named DRI) with one or more associated ORCIDs identifying the data creators. In computer science notation, the DRI tag will have the following structure:

DRI={ORCID1, ORCID2, …}

We note that, implicitly, data consumers are expected to acknowledge/cite any data they use for their scientific work. For the new ethical use of data with DRI, we expect data consumers to follow the approach summarized below and detailed in ***Figure 1***:

1. If DRI present

a. If associated publication(s)

i. Optionally contact data creators
b. If no associated publication

i. Contact data creators
2. Reuse data freely

Sequencing datasets published in public databases (*e.g.*, in Genbank) have tags/attributes/fields that indicate which publications are connected to the respective datasets. Examples of such tags are: i) for GenBank entries - “REFERENCE”, “REFERENCE/AUTHORS”, “REFERENCE/TITLE”, “REFERENCE/JOURNAL”; ii) for

BioSample - “reference for biomaterial”; and iii) for BioProject - “Publications”. The content of these tags is input initially by the data submitter. In theory, they can be updated later either manually by the submitter or automatically by the system. In practice, the submitter seldomly returns to update them or the automatic updating results in completely erroneous references (*e.g.*, https://www.ncbi.nlm.nih.gov/bioproject/PRJEB6121 cites Poole-Wilson & Langer (1975) but should instead cite Probst *et al.* (2014))^18,19^. Most often, missing or erroneous data renders these tags unusable for the purpose of tracking the publication status of the corresponding sequence data. In our roadmap, the DRI, which will contain the ORCID of at least one of the data creators (typically the corresponding creators, *i.e.,* the project leader), will address these issues. The content of this tag would be input/updated solely by the data submitter, preferably during the initial data deposition.

The presence of a DRI tag indicates that data creators prefer to be contacted if a data consumer reuses their data, especially if the respective data have no associated publication. The intentions behind this preference can be manifold, including a willingness to share additional metadata or datasets, or the preference to collaborate to help protect early career researchers’ (*e.g.*, PhD students) ability to finish their studies and graduate. Including one or several ORCID(s) with the DRI will provide a stable point of contact; bolster transparency in science and adherence to the FAIR principles (Findable, Accessible, Interoperable, and Reusable)^20^; and also facilitate science through the exchange of metadata and increased collaboration. There have been instances where such collaboration with data creators, *e.g.*, provision of additional metadata that is not publicly available, has strengthened the content of research studies^22,23^. This should be the norm rather than the exception. In other cases, communication with data creators has allowed proper citation of datasets used, and acknowledgment of funding that supported critical datasets - providing some benefit to the data creator for their data reuse^21,22^. Ideally, in the long run, the DRI tag would help facilitate automatic updates of the publication-associated tags. For example, a Genbank sequence entry could be updated as follows: if the GenBank entry or its corresponding BioProject/BioSamples entries are mentioned in a PubMed-indexed publication together with the ORCID(s) found in the DRI, then the new publication will be automatically added to the respective publication tags. The proposed metadata tag (*i.e.*, the DRI) will substantially reduce the amount of time needed to clarify the status of any public data set, enabling automated rules for dataset screening and reducing tension between data creators and data consumers in the future.

#### Box 3

##### Summary of survey results

A 21-question survey examining the scientific community’s perceptions of data reuse was conducted in January 2024, with the survey distributed through social media, a Nature Microbiology blog, and via email to hundreds of scientists and two dozen scientific societies. A total of 306 respondents contributed to the survey. Responses were summarized for key questions and visualized using Adobe Illustrator. A descriptive analysis of responses to this survey and anonymous raw data to all questions are available in ***Figures S2-10*** and ***Tables S1-5***.

**Figure 2:**
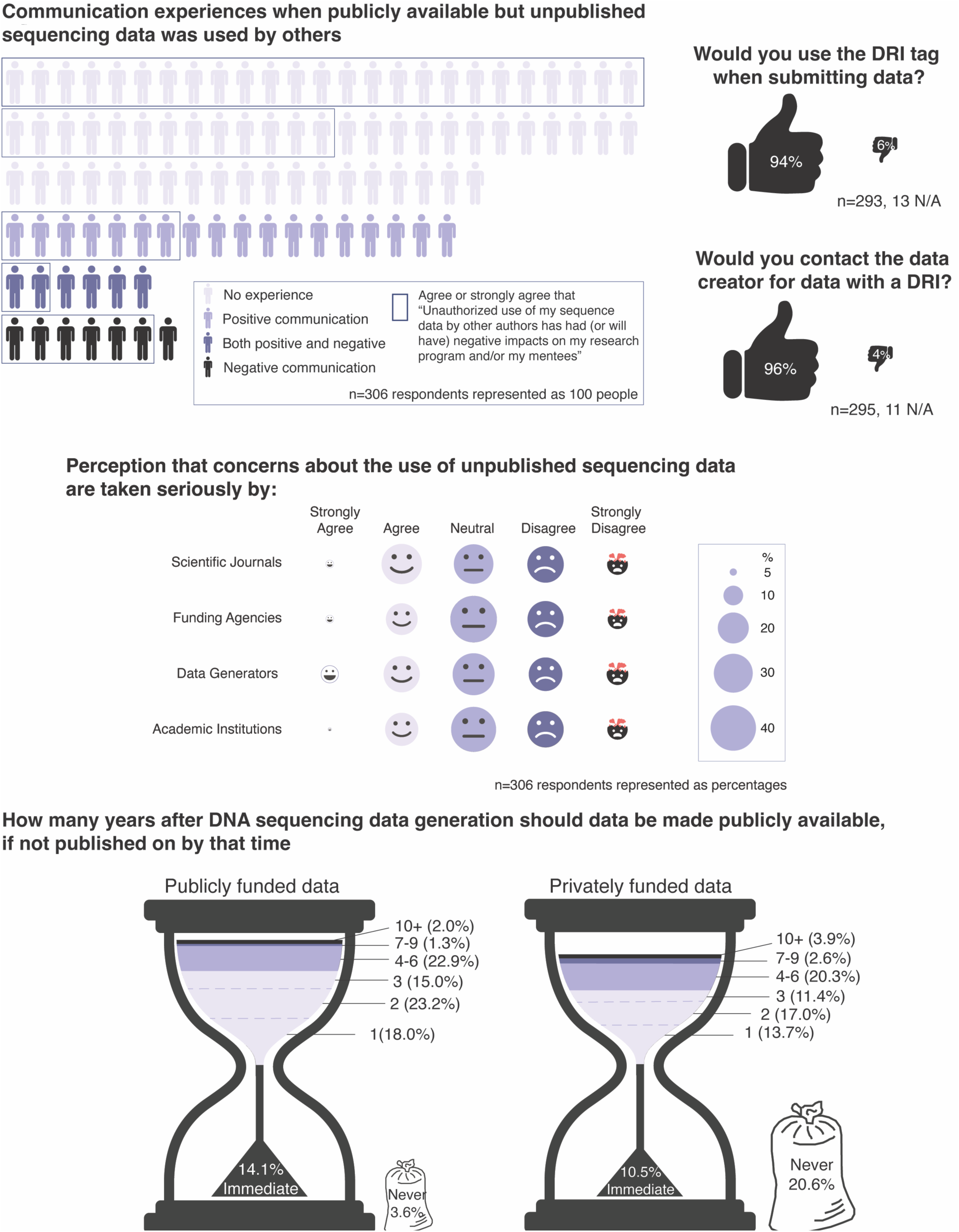
Notes: Positive communication was defined as being contacted before data analysis or publication and asked for collaboration/opinion, or a positive answer when you requested data removal from a manuscript. Negative communication was defined as no contact prior to publication, or refusal to remove data from a manuscript upon request. Respondents selected single year intervals for the data presented in the hourglass image; years 4-6 and 7-9 were combined given very low proportions for all years except year 5. Notably, respondents did not agree on a time interval after which data should be made available in the absence of an available publication (***Figure S7***). Consequently, our roadmap does not include a recommendation as to when publicly available data with a DRI but no publication can be reused without contacting the data creators (see Figure 1).

Ideally, DRIs would propagate to other online databases of -omics sequencing data, like IMG^24^. We note that even tangible incentives (*e.g.*, MG-RAST granting priority access to computing for data made public) have not alleviated data creators’ hesitancy to make their data available^25^. While we do not anticipate the DRI will achieve a complete resolution to this challenge, we do think it is an essential first step in the right direction. The implementation of machine-readable labels reflecting contact information of data creators will permit efficient reuse of data, accelerate scientific discoveries and hopefully lead data creators to more readily share their valuable data sets with the scientific public.

Using the standards devised by the GSC^26^, complete metadata is expected to be made available by data creators. Here, we propose the addition of a DRI tag to GSC metadata. Enabling fair use of public microbiome data will rely on close collaborative work between the data creators, data distributors and data consumers. We envision that the GSC will play a pivotal role in helping all parties (*i.e.*, data distributors) implement a DRI tag within submission systems for microbiome data to public repositories as well as in socializing the new approach to data mining (*i.e.,* the algorithm given above).

### Potential issues and the next steps

The full adoption of a DRI tag for microbiome data in public repositories will ultimately require broad support from the scientific community, data distributors, and from journals and publishing houses. Data creators are encouraged to apply DRI metadata tags to their datasets. This will allow data consumers to connect with data creators to discuss data reuse. We, the Data Reuse Core Team and Data Reuse Consortium, propose that scientific manuscripts partially or exclusively making use of public data should include a written statement by the authors confirming that they have complied with the best practices for public data use. This statement would include protocols for data download and use of analysis tools, or reproducible workflows describing how the tag was incorporated in the workflow or (in case of a missing DRI) how authors adhered to the roadmap outlined in ***Figure 1***.

We are aware that the implementation of the DRI could significantly impact the road from analysis to publication by increasing the workload (*e.g.*, needing to identify email addresses of data creators). This is especially true for projects accessing many datasets (*e.g.*, mining single genes for phylogenetic trees or big data mining projects). However, we are certain that, with time, automated and standardized informatic tools will become available to the entire community to overcome this problem.

At the same time, the high proportion of participants who stated they would respect the DRI when reusing data (266 participants (96.73%); see ***Figure 3***) suggests data creators will be able to more freely and more frequently share their data in the future. This would facilitate science, including data mining, and hugely benefit the scientific community in the long run. Moreover, by fostering collaborations, the scientific best practices and the roadmap for data sharing in microbiome research as introduced here will enable research by lower-resourced laboratories, reducing financial bias in scientific progress. However, we do not expect that the recommendations in this roadmap will be applied retroactively to datasets already deposited in public databases prior to the implementation of the DRI.

**Figure 3:**
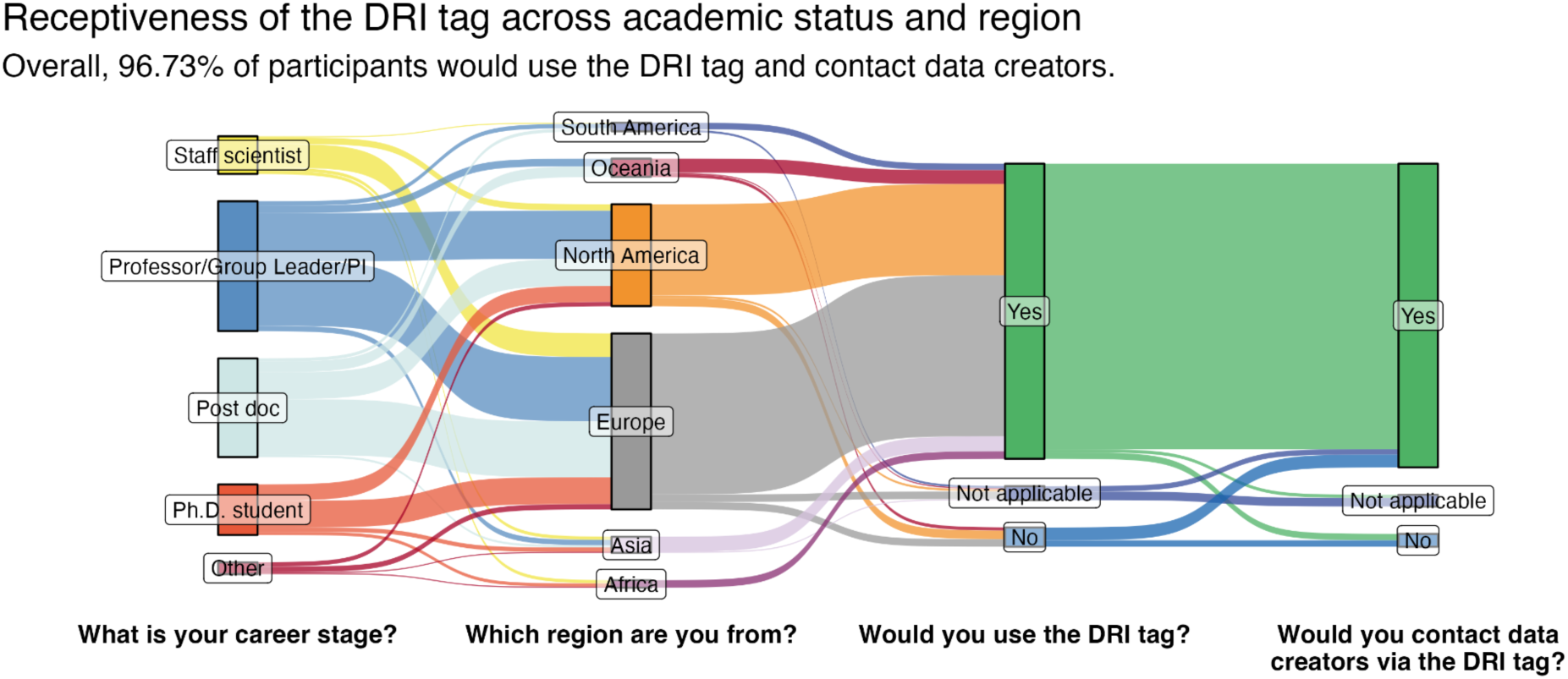
The receptiveness of survey participants towards implementing a DRI tag. Flow height (response category, in color) per stage (survey question, x-axis) is proportional to number of responses. See ***Table S1*** for respondent numbers in each category.

The scale of nucleic acid sequencing data required early pioneers to establish public databases across political borders and to confront data sharing considerations early. Other omics technologies are maturing (*e.g.*, proteomics) such that scientists recognize the need to establish data mining approaches^27^. We propose that this roadmap for fair reuse of public sequencing data should be expanded to other fields including but not limited to proteomics, lipidomics, metabolomics, phenomics, microscopy, and spectroscopy as data mining becomes routine with these types of data.

## Acknowledgements

The Data Reuse Core Team thanks the numerous participants of our survey who chose not to be listed as authors on this manuscript for their valuable perspectives, and the many people who discussed this endeavor with us on multiple occasions. We thank Louisa Rothe for consultations on data visualizations.

## SUPPLEMENTARY INFORMATION

List of scientists supporting this study under the ‘The Data Reuse Consortium’ and respective affiliations:

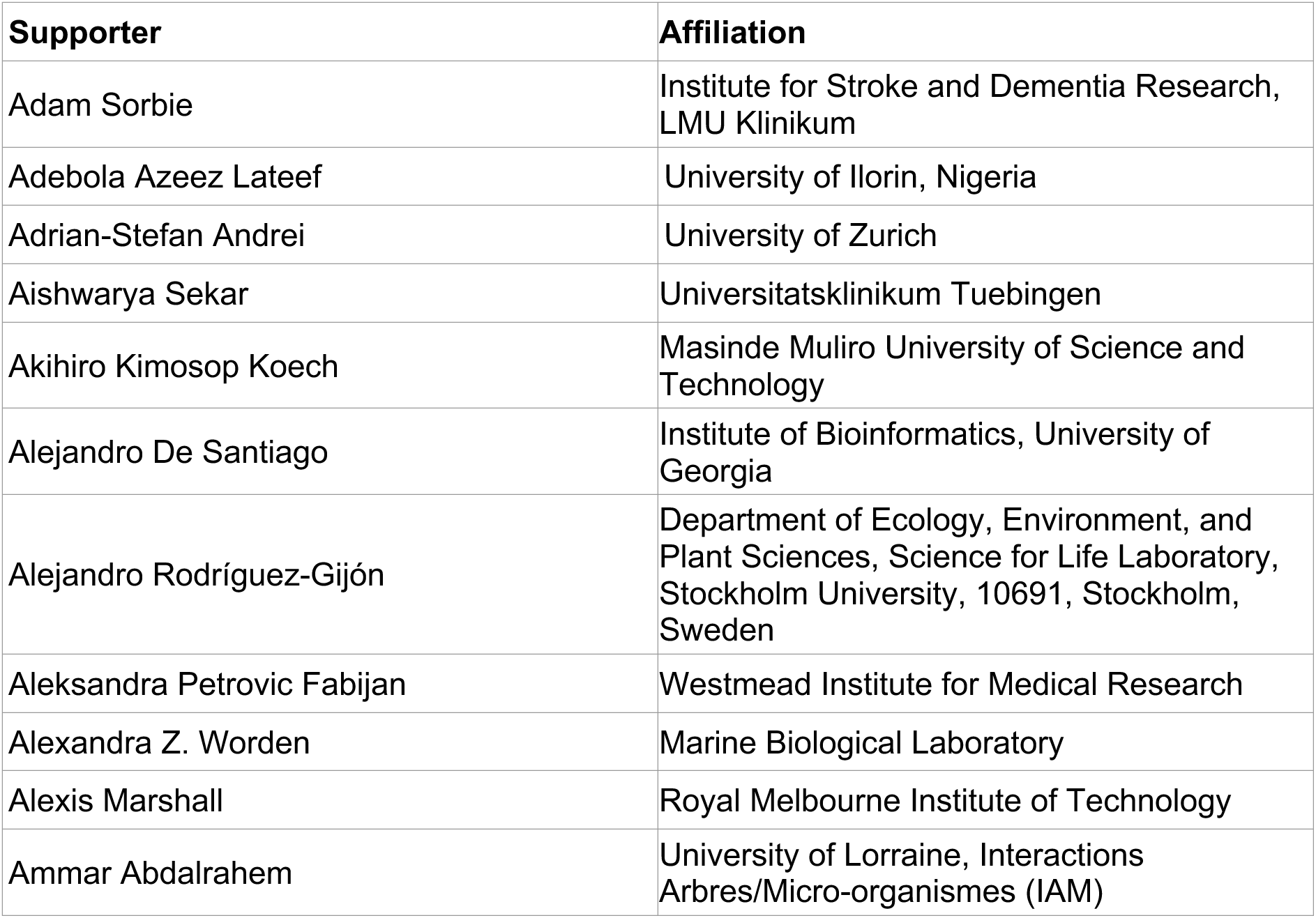
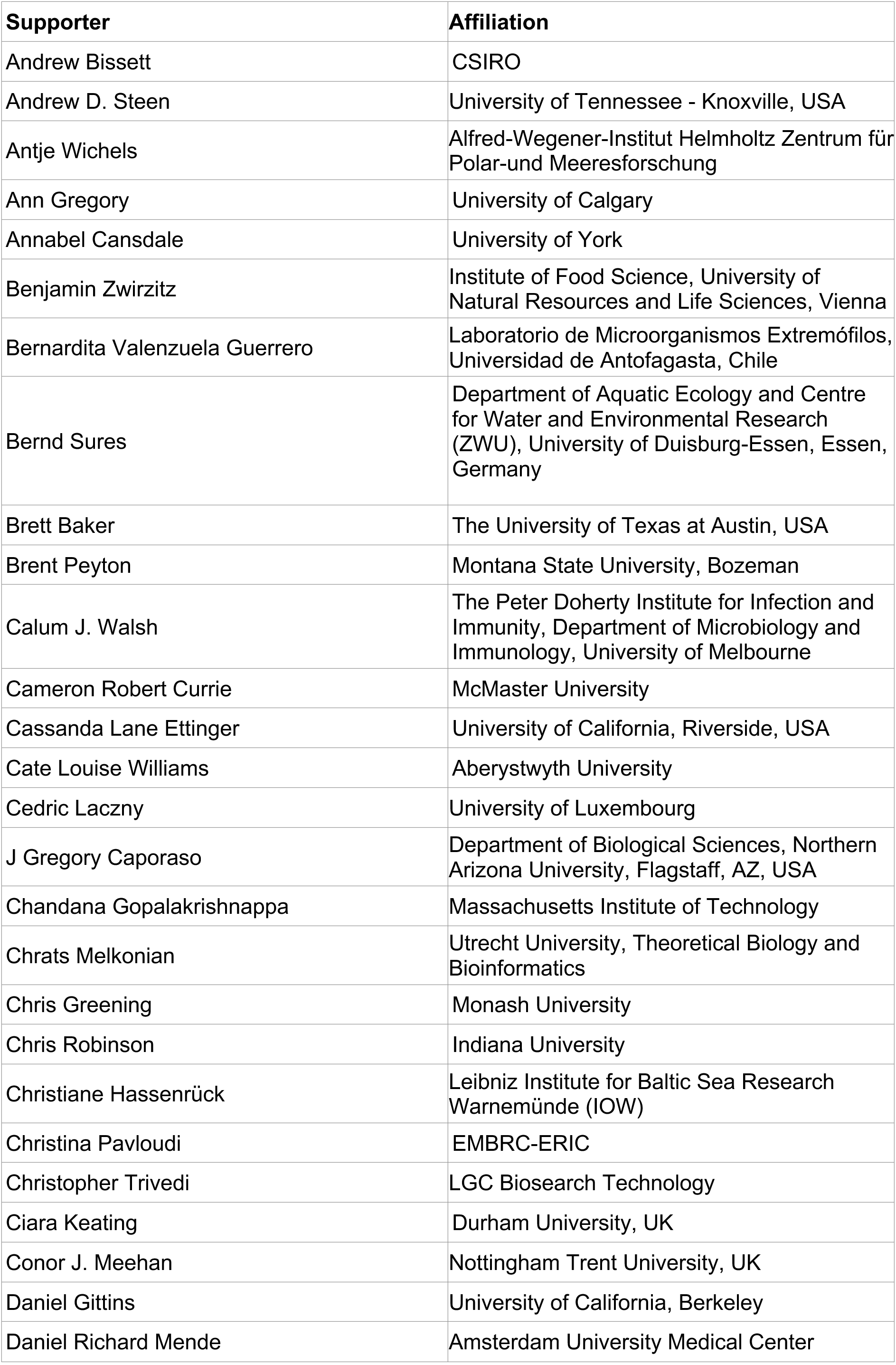
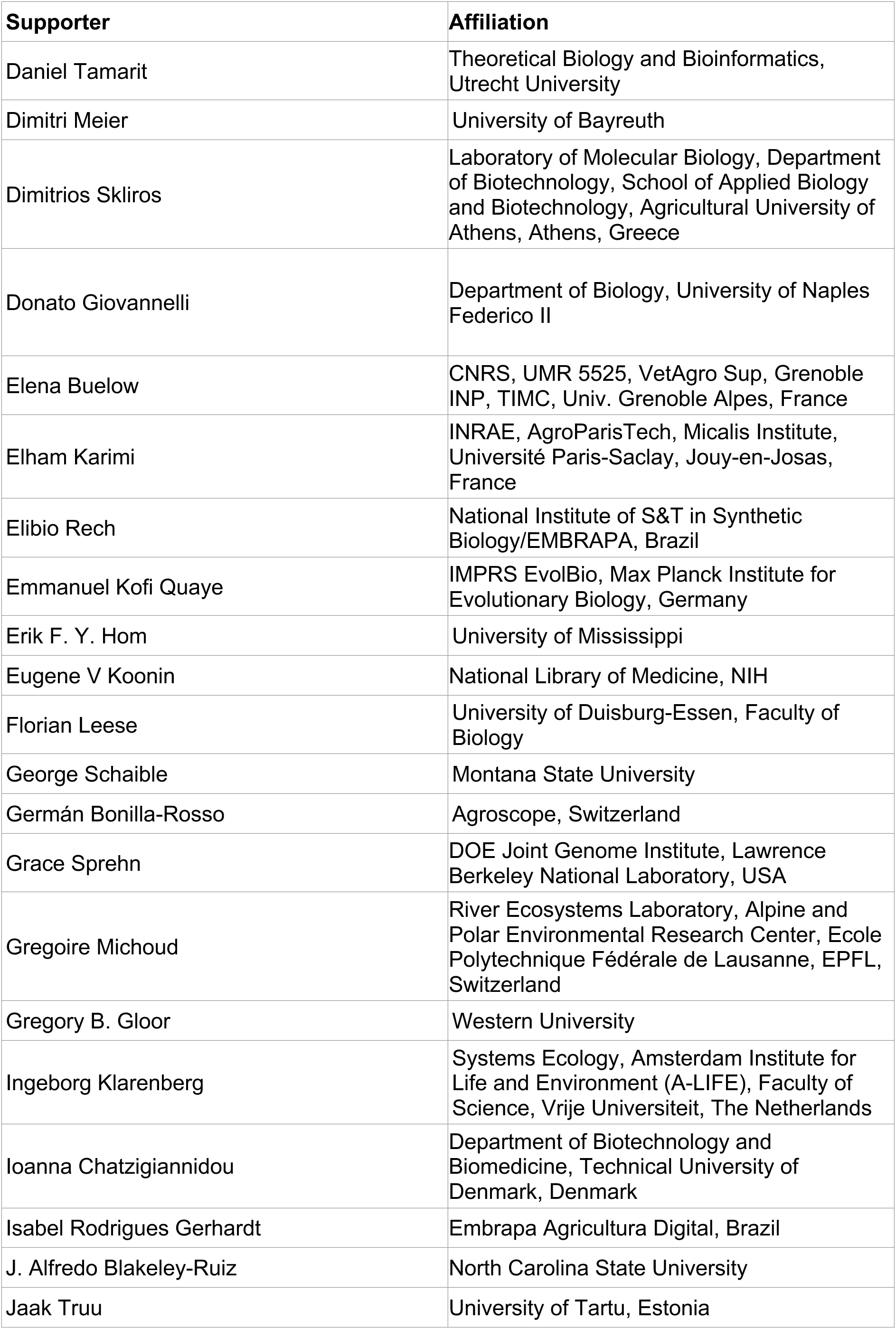
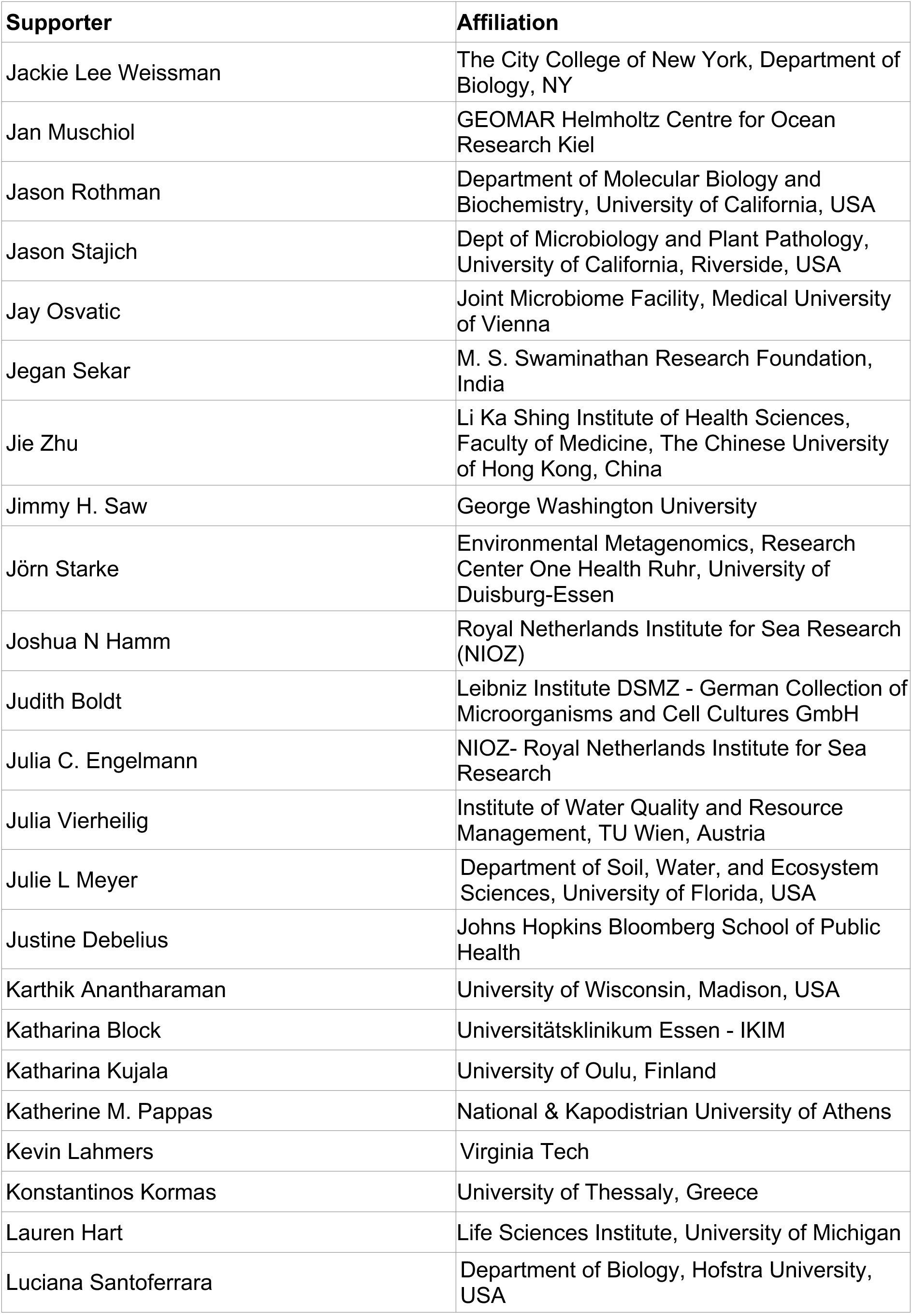
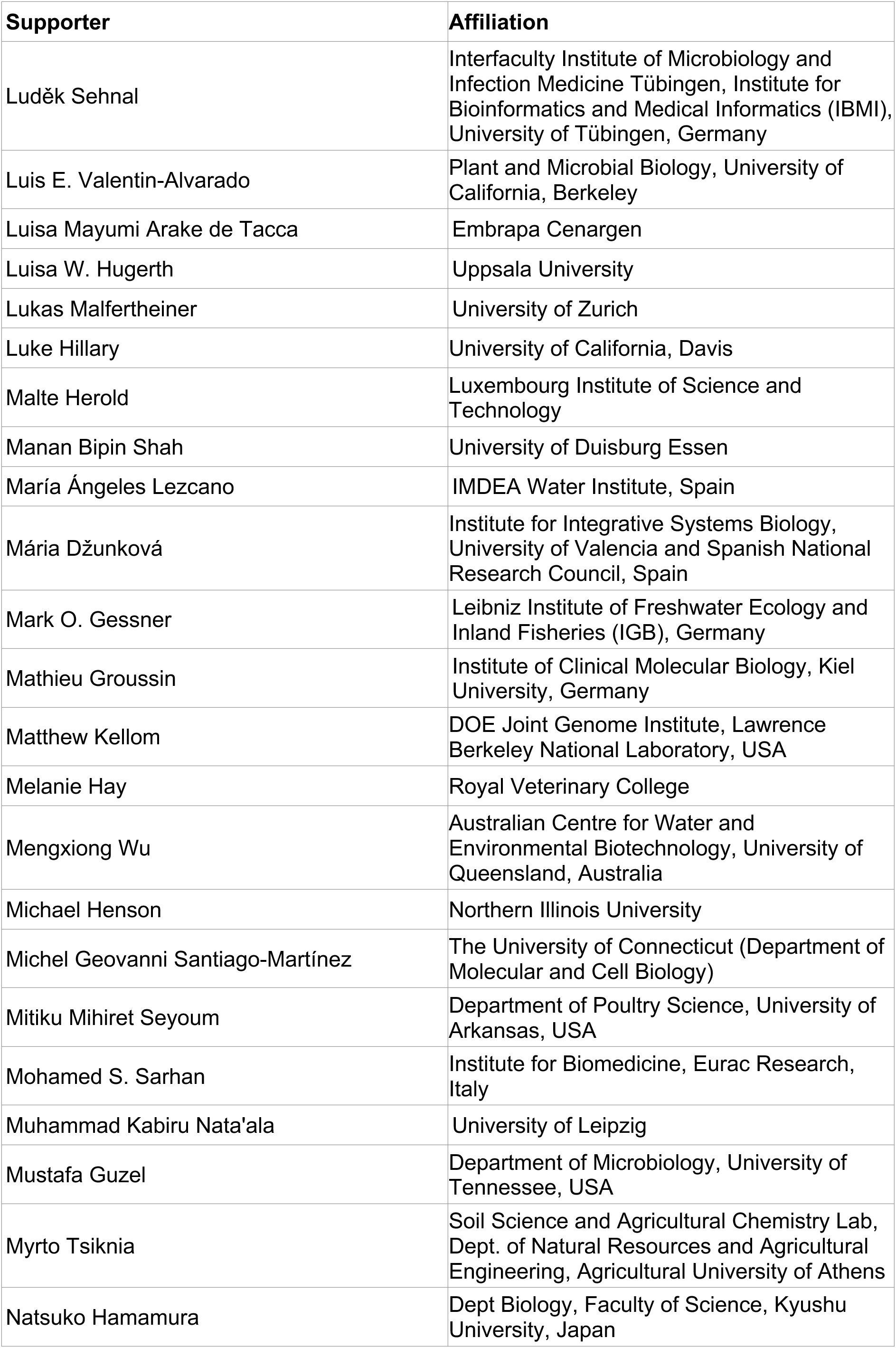
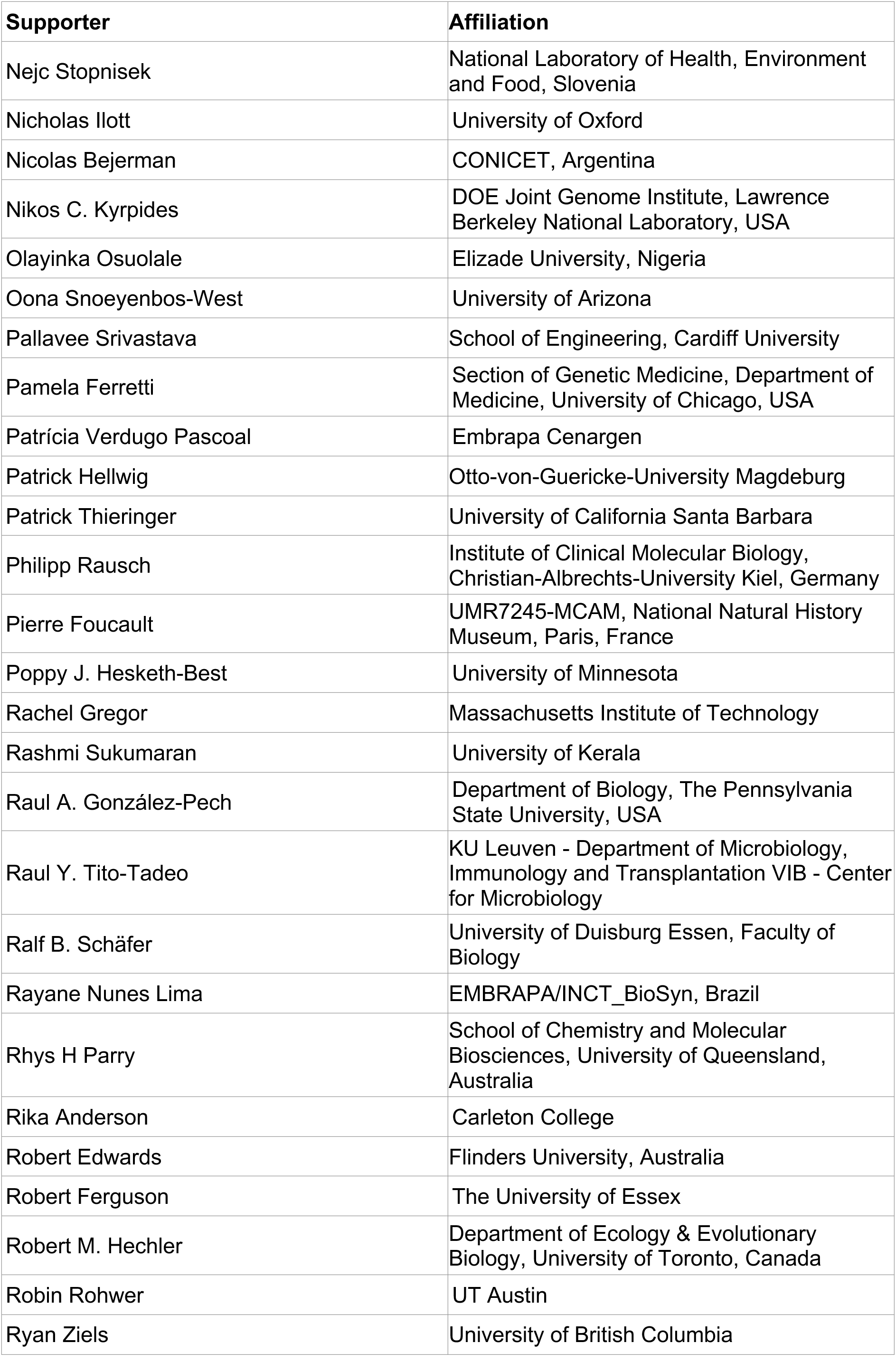
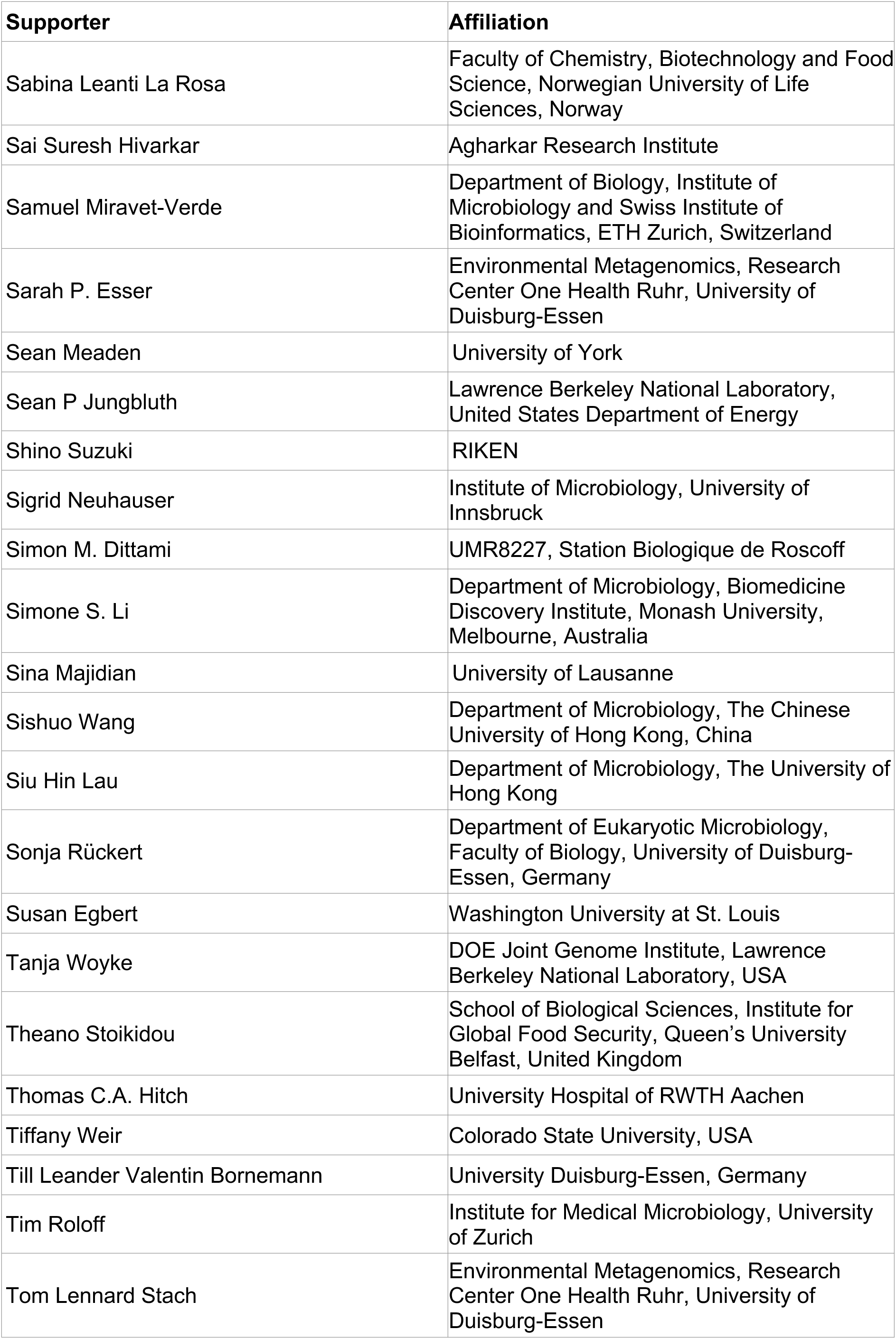
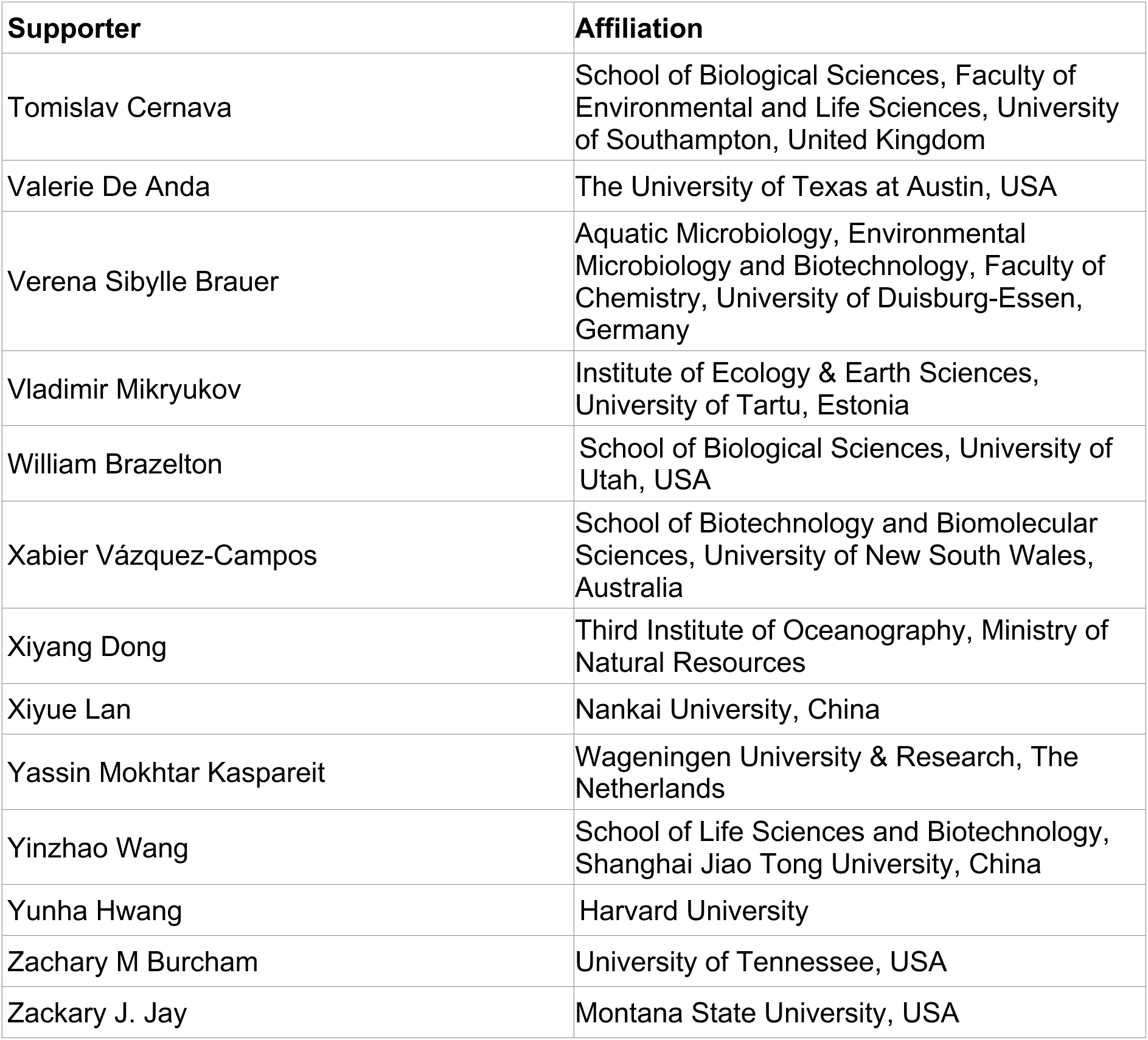

### Methods

Definitions gathered in **Box 1** were created using ChatGPT version 3.5, validated manually via online and literature searches, and edited as needed.

#### Survey of the scientific community

An anonymous survey was created with GoogleForms (https://www.google.com/forms, questions listed below in ‘<u>Questions included in the survey</u>’) and distributed on the 15^th^ of January 2024 to the international scientific community so as to accumulate opinion data related to a number of key topics related to this manuscript. Questions included in the survey were formulated in a neutral fashion with the intent of not biasing responses, which were anonymized to ensure openness and transparency. This survey was online and open for responses over a total of 21 days. Efforts to ensure widespread awareness of the scientific community regarding this survey included actively advertising it within multiple social media platforms (X.com, LinkedIn, Bluesky.app) over the duration of the survey. To achieve this, authors made use of their accounts across these platforms while also leveraging their working group and other institutional accounts. A blog advertising this survey was additionally hosted at the SpringerNature Research Communities blog ( https://communities.springernature.com/posts/contribution-needed-for-developing-a-new-community-standard-for-reusing-sequencing-data). Finally, a total of 78 microbiology institutions from around the world were contacted to increase participation from the global South and underrepresented segments of the global scientific community related to this survey. Raw data pertaining to the anonymous responses to this survey in TSV format has been fully uploaded to the Open Science Foundation (OSF) portal and is available at: https://osf.io/skw4a/?view_only=0c522841fbe24108bc625962a6874a88.

#### Questions included in the survey

1. Which region are you from?
2. What is your career stage?
3. Do you have decision power regarding the release of sequence data to the public for any of the projects you are involved in?
4. Most journals require sequence data associated with a published paper to be made publicly available, e.g., by deposition in public databases. Do you believe that all sequencing data that has been published in a journal (excluding preprint servers) should be publicly available?
5. "Are you aware of the “Bermuda Agreement/Fort Lauderdale Agreement” recommending researchers to make sequence data publicly available within 24 hours after quality control? References: https://web.ornl.gov/sci/techresources/Human_Genome/research/bermuda.shtml#2, https://www.sanger.ac.uk/wp-content/uploads/fortlauderdalereport.pdf
6. In your experience, is the Bermuda Agreement/Fort Lauderdale Agreement commonly adhered to in your country?
7. In your opinion, DNA sequencing data funded by taxpayers’ money should be made publicly available how many years after data generation, if not published on by that time (select “NO” for not having to be released at all, select “0” for immediate release after quality control of raw sequence data, write any number indicating the suggested embargo period in years):
8. In your opinion, DNA sequencing data funded by private funding agencies should be made publicly available how many years after data generation, if not published on by that time (select “NO” for not being released at all, select “0” for immediate release after quality control of raw sequence data, write any number indicating the suggested embargo period in years)
9. In your opinion, concerns about the use of unpublished sequencing data (independent of whether you or another entity created that data) are taken seriously by (select what applies) [Funding agencies]
10. In your opinion, concerns about the use of unpublished sequencing data (independent of whether you or another entity created that data) are taken seriously by (select what applies) [Scientific journals]
11. In your opinion, concerns about the use of unpublished sequencing data (independent of whether you or another entity created that data) are taken seriously by (select what applies) [Academic institutions]
12. In your opinion, concerns about the use of unpublished sequencing data (independent of whether you or another entity created that data) are taken seriously by (select what applies) [Data generators (e.g., taxpayer-funded sequencing facilities).]
13. Please indicate your agreement with the following statement: “I am concerned about other people publishing on my sequence data before I have published on them”:
14. Has your publicly available but unpublished sequencing data been used by others in the past?
15. Have you ever had a positive communication experience when your publicly available but unpublished sequencing data was used by others in the past? (positive communication is defined here as being contacted before data analysis or publication and asked for collaboration/opinion, or a positive answer when you requested data removal from a manuscript)
16. Have you ever had a negative communication experience when your publicly available but unpublished sequencing data was used by others in the past? (negative communication defined here as no contact prior to publication, or refusal to remove data from a manuscript upon request)
17. Please indicate your agreement with the following statement: “Unauthorized use of my sequence data by other authors has had (or will have) negative impacts on my research program and/or my mentees.”
18. Would you define yourself as a (for definitions, please see the introductory paragraph):

a. You have responded "Other" to the previous question: How do you define yourself?
19. Would you use the DRI tag when submitting your data to a public database to facilitate collaborative reuse of your data?
20. If a dataset included a DRI tag, indicating the data creator prefers to be contacted prior to reuse of their data, would you respect this and contact them?
21. Would you like to participate in this manuscript as a co-author as part of the “Data Reuse Consortium”? Please note that the information you just provided will remain anonymous whichever option is chosen. If you agree, you will need to provide your full name and contact email. All contact information will be kept a second, independent database, that cannot be linked to this survey.

#### Data analysis of survey data

Raw data for this survey was imported into R 4.3.1 running in RStudio 2023.12.0 (Build 369) using the tidyverse ecosystem of data analysis packages to read TSV inputs (*readr*), filter

(*dplyr*) and generate plots (*ggplot2*)^1,2^. Sankey diagrams were generated via the *ggsankey* package (https://github.com/davidsjoberg/ggsankey). Tables were generated with the *kableExtra* package^3^. R code as well as system and package versions used for all data analysis in this manuscript are publicly available at a GitHub repository: https://github.com/GeoMicroSoares/DataUsage_Data_Analysis. Data visualization for the **Box 3** infographic was conducted in Adobe Illustrator using copyright-free images. Proportional scaling was applied for size comparisons. The emoji bubble plot was generated in R using *ggplot2* and standard bubbles replaced with scaled emoji icons in Adobe Illustrator.

### Results

#### Growth of SRA sequencing data since 2007 and growth prediction in the future

**Figure S1:**
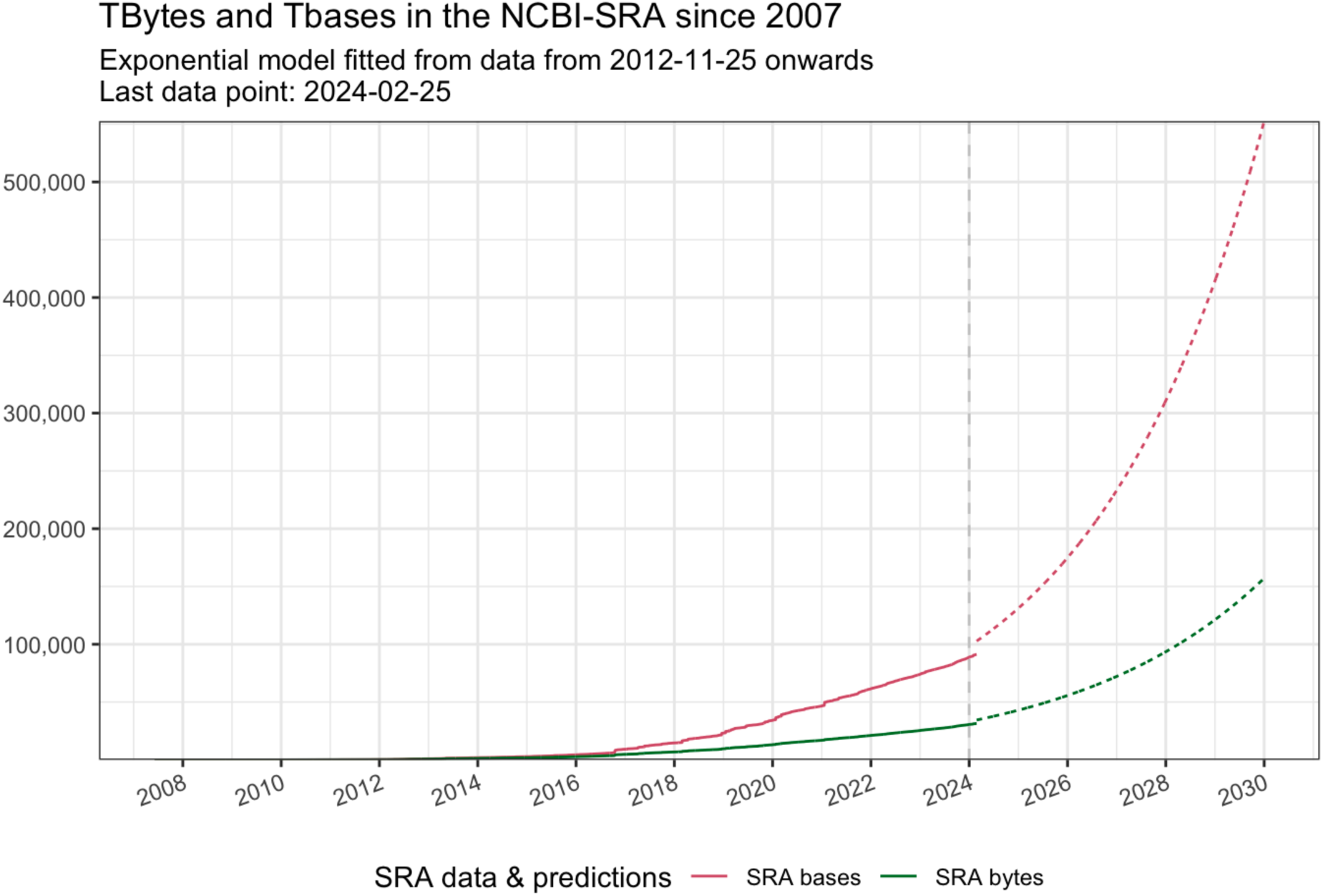
Total bases and bytes (color) hosted by SRA (y-axis) over time (x-axis). Dashed lines correspond to a fitted exponential model using data from 2012-11-25 onwards. Vertical dashed gray line indicates 2024-01-01.

#### Descriptive analysis of the scientific community survey

**Figure S2:**
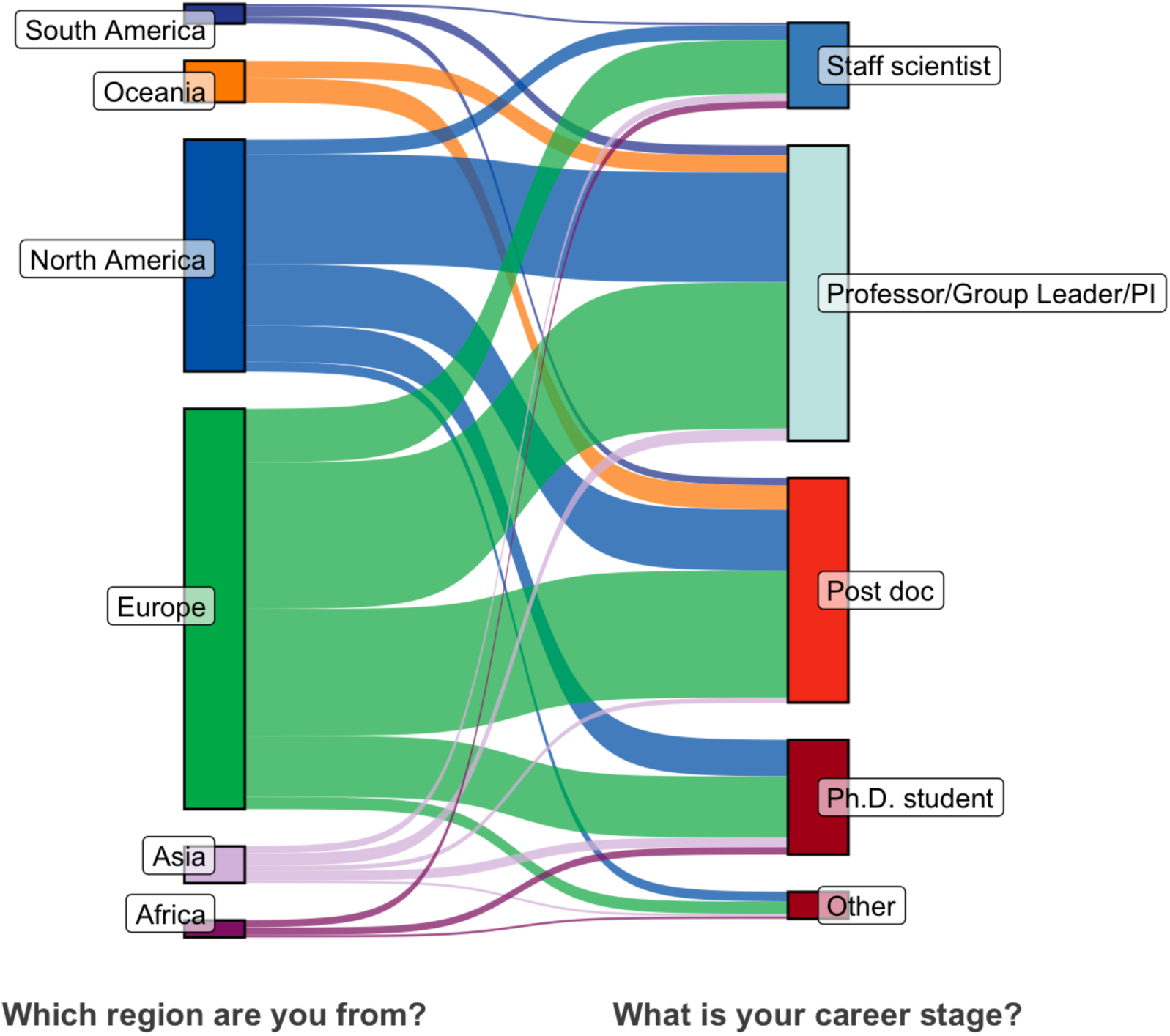
Sankey diagram depicting relationships between country of origin and career stage. Node width is proportional to number of data points (responses).

**Table S1:** Summary table depicting numbers and percentages of participant responses by career stage and country of origin. Cells containing “No answers” relate to conditions for which no responses were obtained.

| Which region are you from? | Professor/Group Leader/PI | Post doc | Staff scientist | Ph.D. student | Other |  |
| --- | --- | --- | --- | --- | --- | --- |
| Africa | No answers | No answers | 1% (3) | 1% (3) | 0.3% (1) | 2.3% (7) |
| Asia | 1.6% (5) | 0.7% (2) | 1% (3) | 1.3% (4) | 0.3% (1) | 4.9% (15) |
| Europe | 19.6% (60) | 17% (52) | 7.2% (22) | 8.2% (25) | 1.6% (5) | 53.6% (164) |
| North America | 14.7% (45) | 8.2% (25) | 2% (6) | 4.9% (15) | 1.3% (4) | 31% (95) |
| Oceania | 2.3% (7) | 3.3% (10) | No answers | No answers | No answers | 5.6% (17) |
| South America | 1.3% (4) | 1% (3) | 0.3% (1) | No answers | No answers | 2.6% (8) |
|  | 39.5% (121) | 30.1% (92) | 11.4% (35) | 15.4% (47) | 3.6% (11) | 100% (306) |

**Figure S3:**
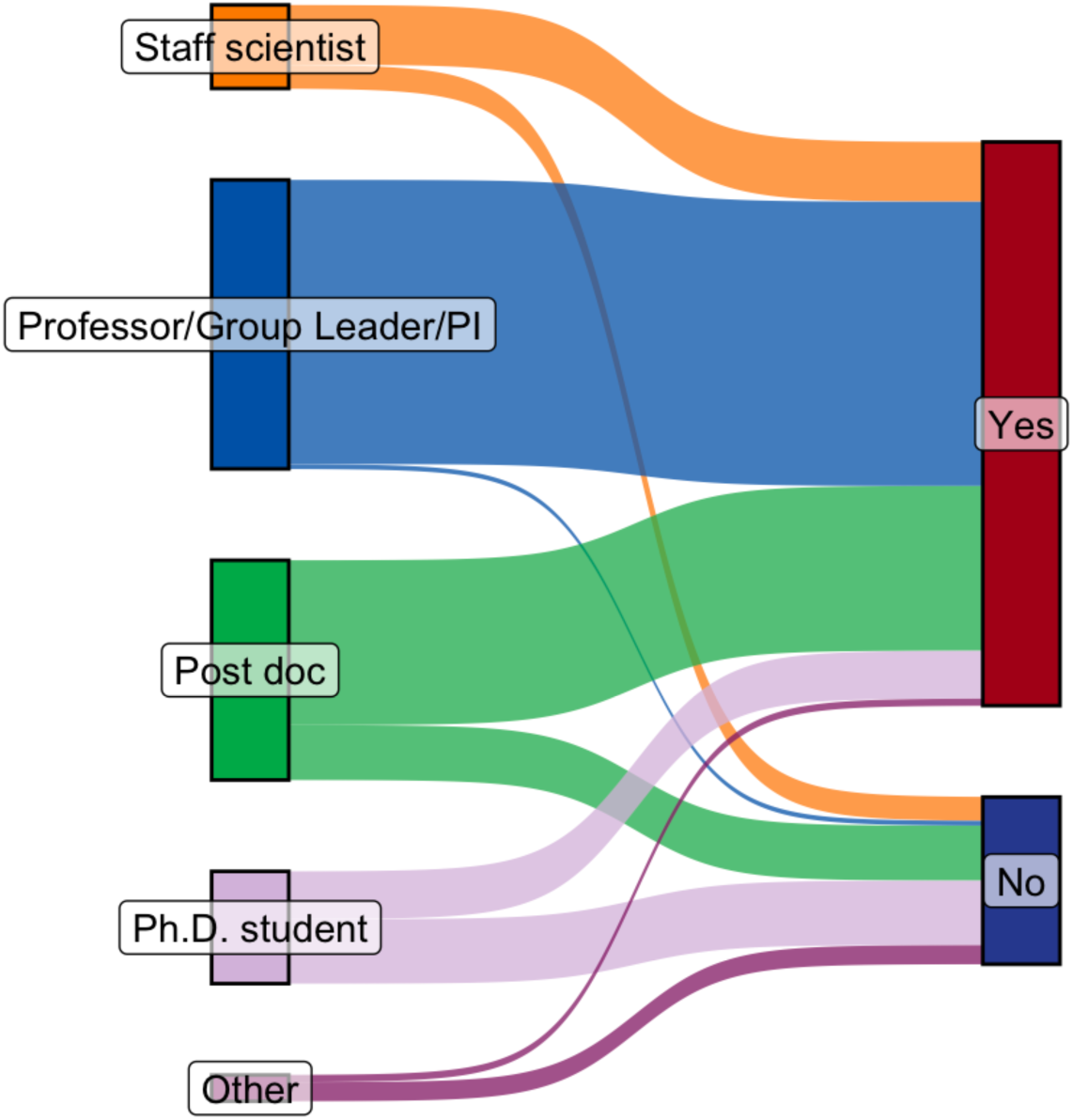
Sankey diagram depicting relationships between career stage and decision power for release of sequence data to the public (exact numbers and percentages in **Table S2**). Node width is proportional to the number of data points (responses).

**Table S2:** Summary table depicting numbers and percentages of participant responses regarding career stage and decision power for release of sequence data to the public.

| Career stage | Yes | No |
| --- | --- | --- |
| Other | 1% (3) | 2.6% (8) |
| Ph.D. student | 6.5% (20) | 8.8% (27) |
| Post doc | 22.5% (69) | 7.5% (23) |
| Professor/Group Leader/PI | 38.9% (119) | 0.7% (2) |
| Staff scientist | 8.2% (25) | 3.3% (10) |

**Table S3:** Summary table depicting numbers and percentages of participant responses by career stage and their opinion regarding public availability of published sequencing data.

| Career stage | Yes | No |
| --- | --- | --- |
| Other | 3.3% (10) | 0.3% (1) |
| Ph.D. student | 14.7% (45) | 0.7% (2) |
| Post doc | 29.4% (90) | 0.7% (2) |
| Professor/Group Leader/PI | 36.9% (113) | 2.6% (8) |
| Staff scientist | 10.5% (32) | 1% (3) |

**Figure S4:**
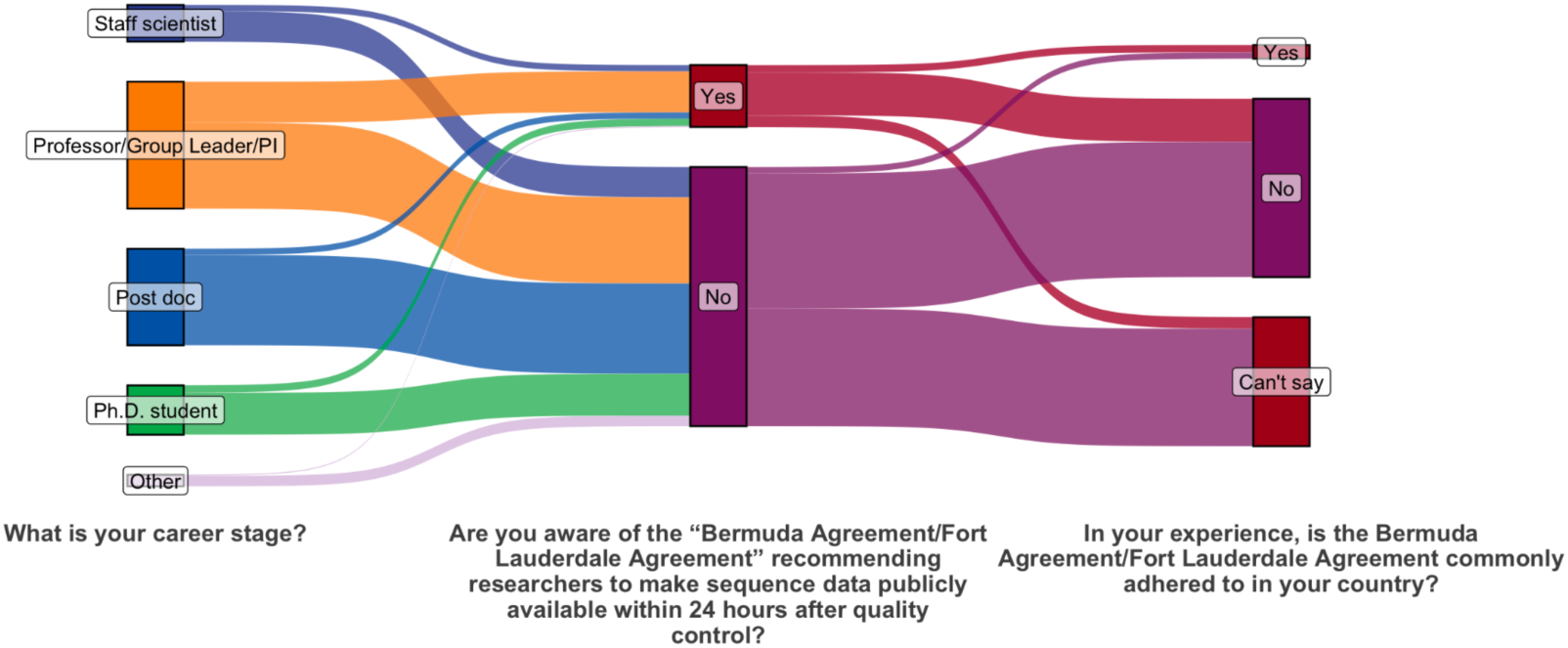
Sankey diagram depicting relationships between career stage and awareness of the existence of the Bermuda Agreement and its implementation in their countries. Node width is proportional to the number of data points (responses).

**Figure S5:**
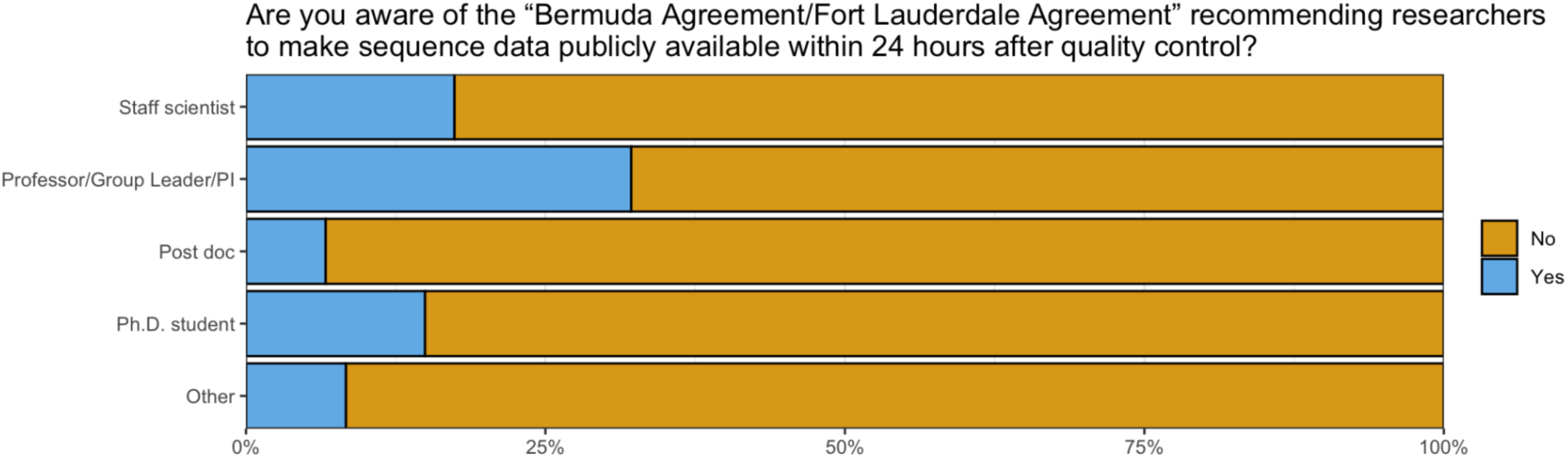
Barplot depicting data trends supporting **Figure S4** - y-axis depicts participant career stage, x-axis percentages and bar color indicates response. Plot title indicates the question to which this subset of the data pertains to.

**Figure S6:**
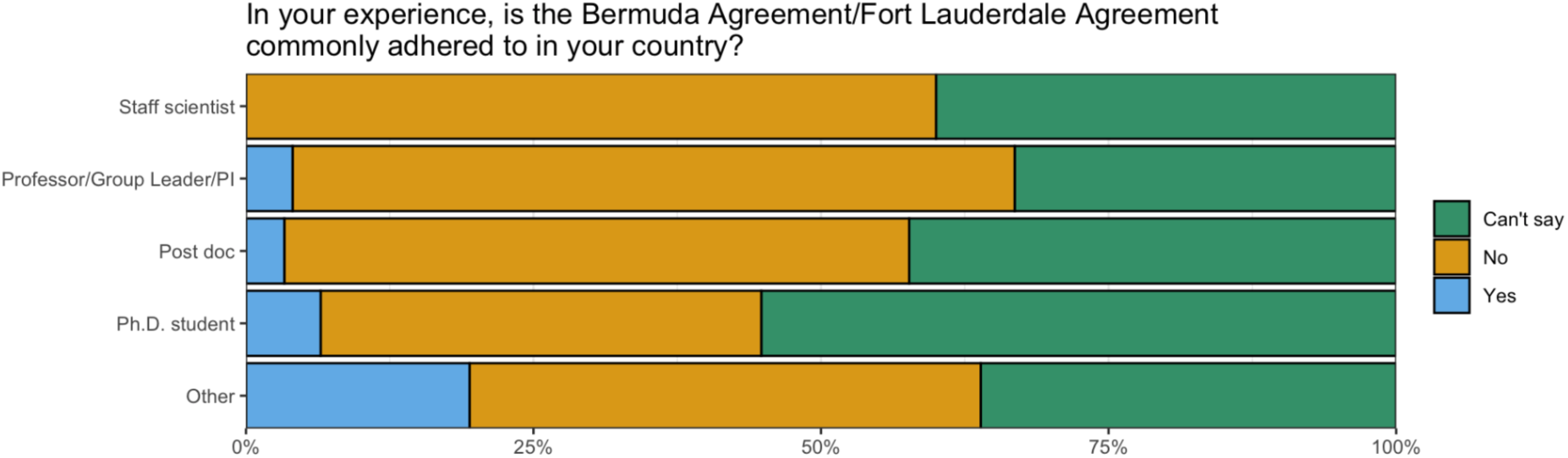
Barplot depicting data trends supporting **Figure S4** - y-axis depicts participant career stage, x-axis percentages and bar color indicates response. Plot title indicates the question to which this subset of the data pertains to.

**Figure S7:**
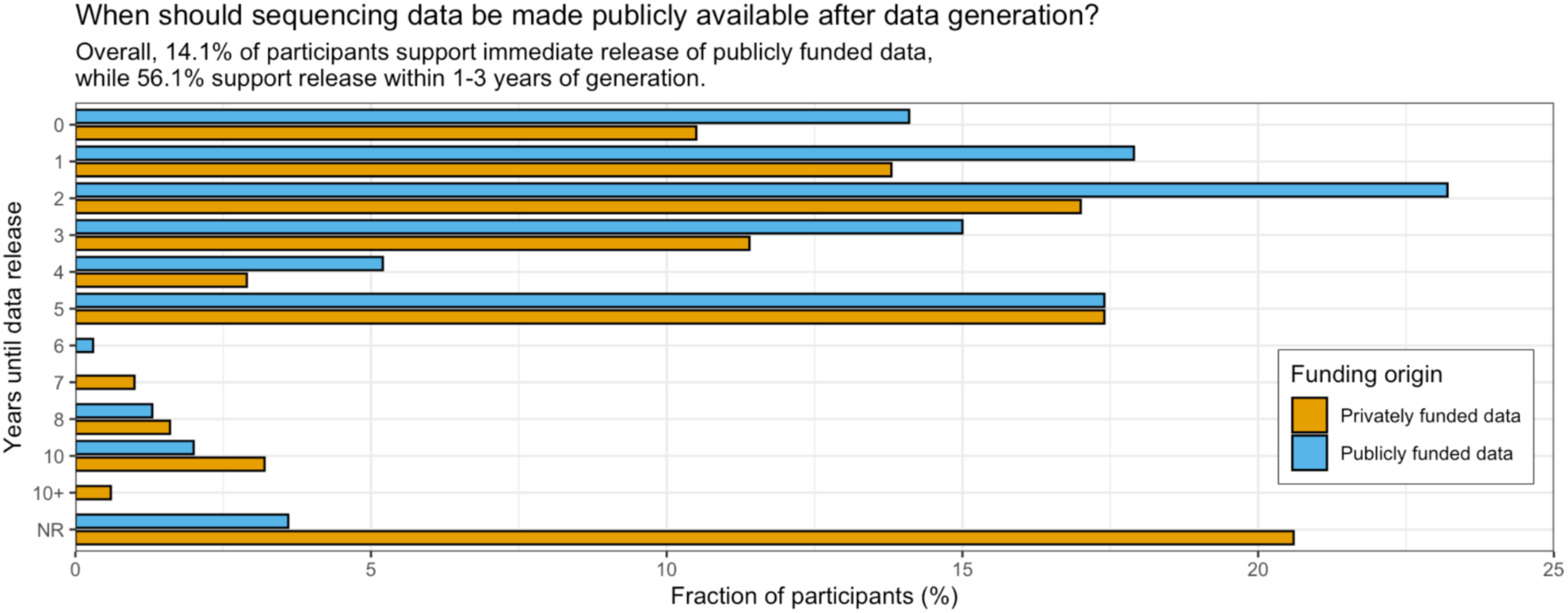
Grouped barplot describing participant support (in percentage - x-axis) for time frames (y-axis) to make sequencing data publicly available given the origin of the funding used to generate such data (color). NR indicates ‘never releasing’.

**Figure S8:**
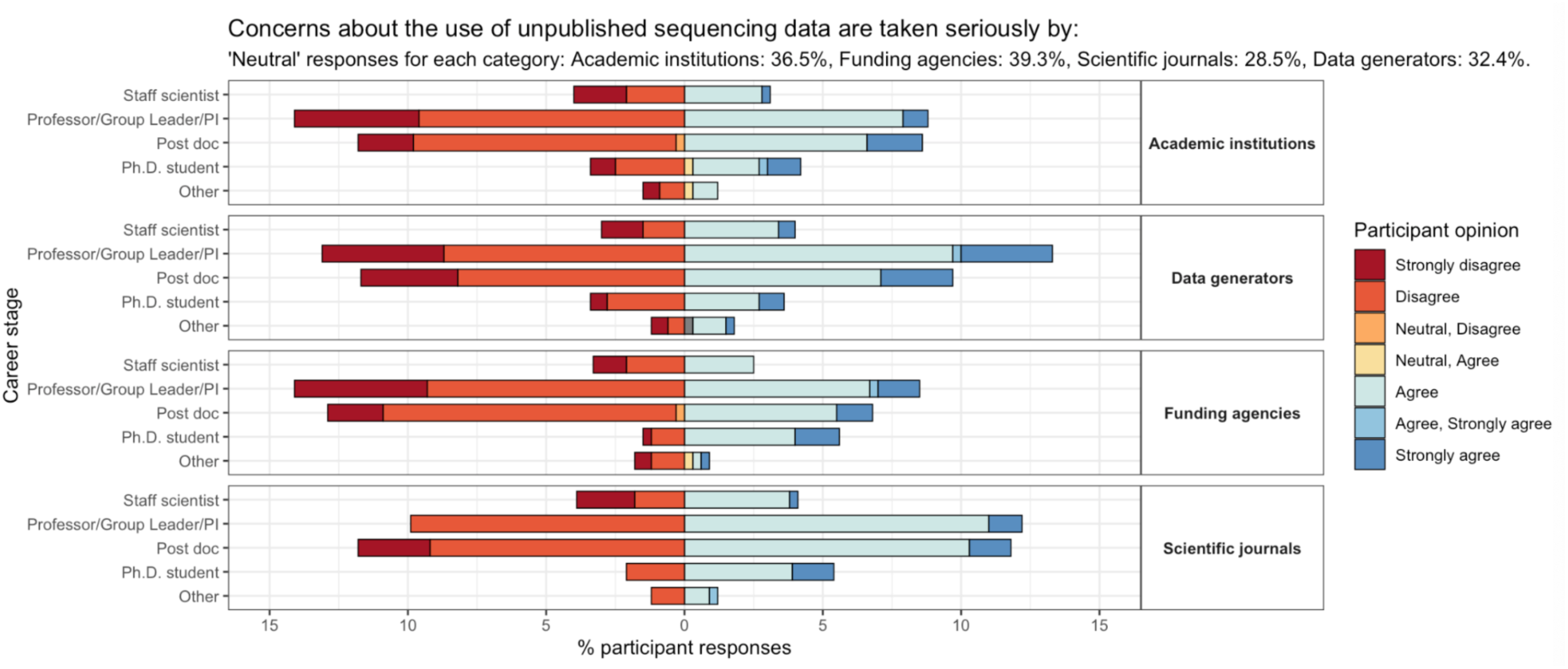
Barplot describing participant opinion (color, x-axis depicts percentages) regarding how seriously their concerns regarding unpublished sequencing data are taken by academic institutions, data generators, funding agencies and scientific journals (vertical facets) across career stage (y-axis).

**Figure S9:**
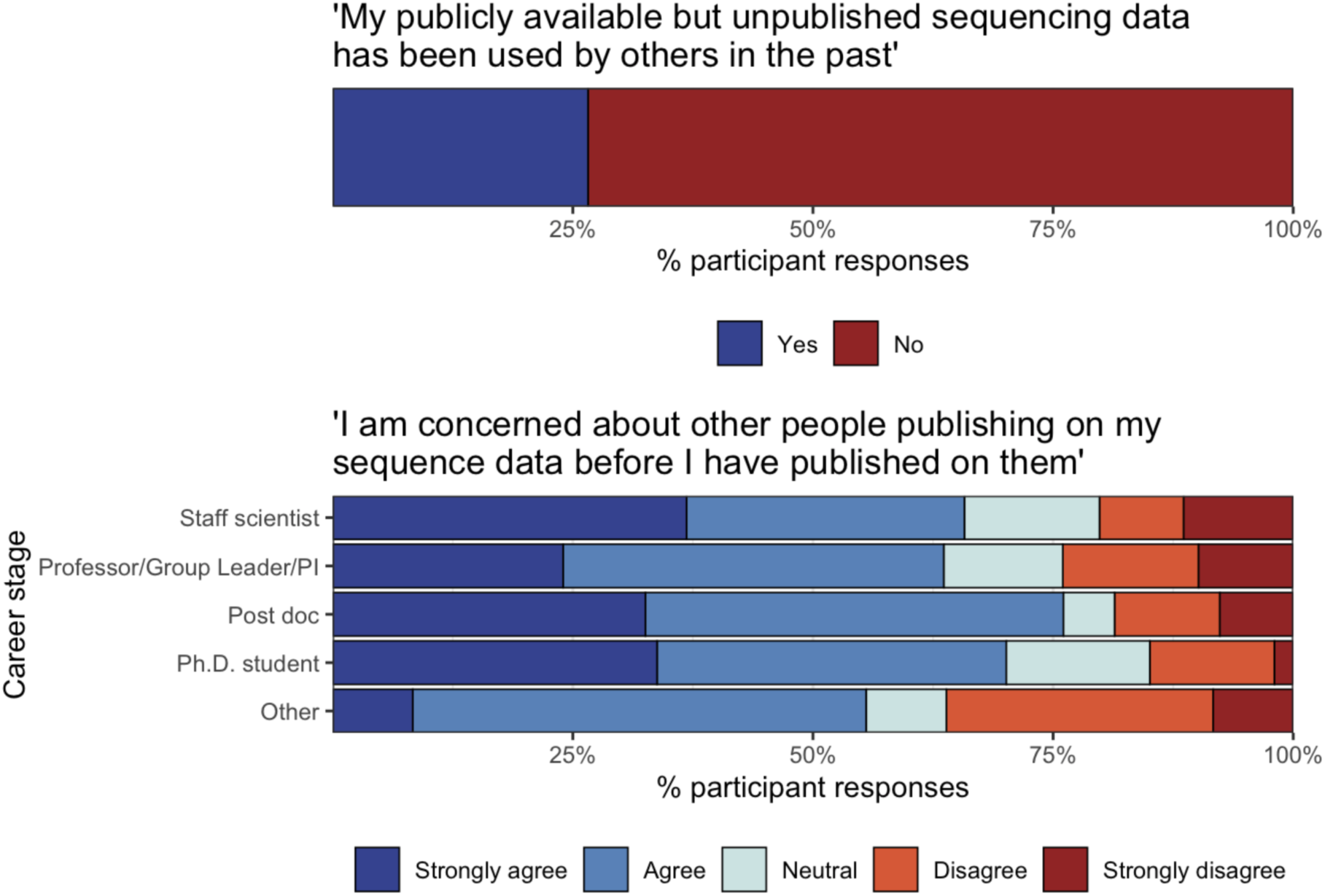
Barplots related to participant opinions on use of their publicly available but unpublished sequencing data (see plot titles). In the top plot, ‘Yes’ or ‘No’ (color) relate to the percentage (x-axis) of participants agreeing or not with the statement in the plot title. The bottom plot depicts agreement or disagreement (color) in percentage (x-axis) with the statement on the plot title across career stages (y-axis).

**Figure S10:**
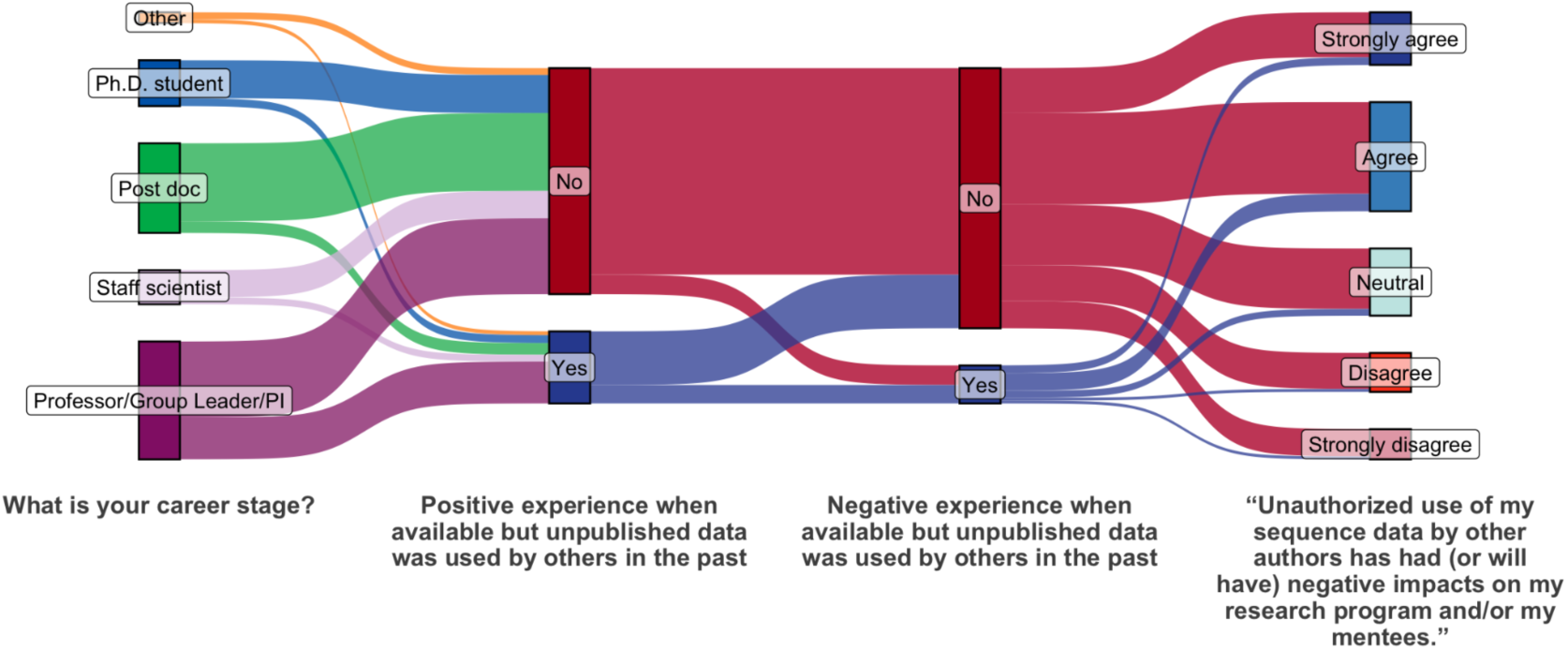
Sankey diagram depicting relationships between career stage and experience quality upon use by others of unpublished data in the past and its consequences (questions in the x-axis). Node width is proportional to the number of data points (responses).

**Table S4:** Summary table depicting numbers and percentages of participant responses regarding the consequences of sequence data usage by others in the past and its impacts (supportive of **Figure S9**).

| “Unauthorized use of my sequence data by other authors has had (or will have) negative impacts on my research program and/or my mentees.” | Count | Fraction (%) |
| --- | --- | --- |
| Strongly agree | 54 | 17.6 |
| Agree | 112 | 36.6 |
| Neutral | 69 | 22.5 |
| Disagree | 40 | 13.1 |
| Strongly disagree | 31 | 10.1 |

**Table S5:** Summary table depicting numbers and percentages of participant responses given their definition as data consumer, creator, both or other definitions.

| Would you define yourself as a: | Count | Fraction (%) |
| --- | --- | --- |
| **Both the above** | 241 | 78.8 |
| **Data consumer** | 16 | 5.2 |
| **Data creator** | 47 | 15.4 |
| **Other** | 2 | 0.7 |

